# Decoding the restriction of T cell receptors to human leukocyte antigen alleles using statistical learning

**DOI:** 10.1101/2025.02.06.636910

**Authors:** Hesham ElAbd, Aya K.H. Mahdy, Eike Matthias Wacker, Maria Gretsova, David Ellinghaus, Andre Franke

## Abstract

Conventional T cells recognize peptides presented by the human leukocyte antigen (HLA) proteins through their T cell receptors (TCRs). Given that thousands of HLA proteins have been discovered, each presenting thousands of different peptide antigens, decoding the cognate HLA protein of a TCR experimentally is a challenging task. To address this problem, we combined statistical learning methods with a unique dataset of paired T cell repertoires and HLA allele information for 6,794 individuals. This enabled us to discover 53,870 T cell receptor alpha (TRA) and 1,095,576 beta (TRB) clonotypes that were associated with 437 unique HLA alleles. The identified clonotypes were targeting prevalent infections, *e.g.* influenza, cytomegalovirus and Epstein-Barr virus. Utilizing these clonotypes, we developed statistical models that impute the carriership of common HLA alleles from the TRA- or TRB-repertoire. In conclusion, the identified allele-associated clonotypes encode the HLA fingerprints and the immune exposure history of individuals and populations.

## Introduction

T cells are an essential component in fighting infections and controlling cancers^1^ as well as mediating tissue homeostasis^2^. T cells are characterized by a high diversity in their receptors, forming the basis of their ability to recognize a broad spectrum of antigens. Based on the nature of the antigens they recognize, T cells are divided into two main categories: conventional T cells which recognize peptides presented by the <u>h</u>uman leukocyte <u>a</u>ntigen (HLA) proteins and unconventional T cells which recognize a wide array of non-peptide antigens presented by non-HLA molecules such as vitamin B metabolites presented by the MR1 protein^3^, and lipids presented by the CD1 molecules^4^. T cell receptors (TCRs) are heteromeric proteins that are made from two antigen-binding chains. In humans, there are four antigen binding chains, namely α, β, γ, and δ, where α and β preferentially dimerize together to form αβ TCRs, γ and δ chain bind together to form γδ TCRs. Conventional T cells rely on the αβ TCRs while unconventional T cells utilize both αβ TCRs such as mucosal-associated invariant T (MAIT) cells which recognize MR1-presented antigens^3^ and γδ TCRs such as Vγ9Vδ2 T cells which recognize phosphoantigens on butyrophilins^5^. We here focused on conventional αβ TCRs.

Each α and β chain is generated through a somatic recombination process termed V(D)J recombination in which the recombination between different V genes, D genes (only in the β chains), and J genes coupled with the random insertion and deletion of nucleotides generates diverse antigen binding chains. Recent advances in next-generation sequencing have advanced our ability to profile the collection of TCR antigen binding chains present in a sample, *i.e.* the α (TRA) or the β (TRB) immune repertoire. Bulk TCR-Seq is a commonly used method to profile the immune repertoire by identifying the collection of the V(D)J recombination events constituting the repertoire^6^. Although TCR-Seq can be conducted using different assays and methods, it is commonly performed using a multiplex PCR reaction that amplifies the DNA-encoded V(D)J recombination products followed by deep sequencing and bioinformatics analyses to identify and quantify the expansion of different clonotypes in the repertoire of a given sample^6^.

Given that different studies have shown that HLA alleles have a strong influence on the TCR repertoire^7,8^, it was a logical consequence that subsequent studies aimed at associating T cell clonotypes with different HLA alleles. One of the earliest studies was performed by Emerson and colleagues who used a statistical framework to analyze the TRB repertoire of 666 individuals with matched HLA-A and HLA-B allele-calls, enabling them to identify thousands of TRB clonotypes that were restricted to multiple HLA-A and HLA-B alleles^9^. This was followed by Ortega *et al.*^10^ who analyzed multiple publicly available datasets of paired HLA genotyping and TCR repertoires to develop models for imputing HLA alleles from the TCR repertoire. Lastly, in a pre-print manuscript, Zahid *et al.*^11^ described the analysis of paired TRB and HLA genotypes for ∼4,000 samples and the development of a machine-learning framework for predicting the HLA background of a sample using its TRB repertoire.

These studies have established the feasibility of performing HLA typing from the T cell repertoire as well as the utility of large-scale statistical analyses in identifying public clonotypes associated with different HLA alleles. However, all mentioned studies did not release a publicly available tool for imputing HLA allele from TCR repertoires, neither did they provide a detailed investigation of the factors shaping the predictive performance of these models. Therefore, a publicly available dataset of TRA and TRB clonotypes that are restricted to common HLA proteins remains unavailable. Such a dataset would aid in the analysis of TCR-Seq studies by identifying the cognate HLA alleles to common clonotypes. To address these limitations, we assembled a large dataset of 1,240 TRA and 5,554 TRB repertoires with their HLA information, enabling us to identify the cognate HLA alleles for 53,870 TRA and 1,095,576 TRB clonotypes. Subsequently, we used the identified alleles-associated clonotypes to build an HLA imputation framework based on these clonotypes.

## Results

### Assembly of a large dataset of paired T cell repertoires and HLA genotypes

Given that sample size significantly influences the performance of the imputation models^10^, we set out to generate and assemble a large dataset of paired TCR repertoires and HLA genotypes (**Table 1**). The TCR repertoires were generated using two TCR-Seq methods, namely, Adaptive Biotechnologies ImmunoSEQ assay (Seattle, USA) and αβ TCR profiling kits from MiLaboratories (Sunnyvale, California), while HLA genotypes were imputed from dense SNP genotyping arrays^12^ (**Online Methods**). This enabled us to build, to the best of our knowledge, the largest dataset of paired T cell repertoires with HLA genotypes covering 433 unique HLA alleles across three different populations at the time of writing (Germany, USA and Norway). Subsequently, we split this dataset into three parts. First, paired TRB-HLA alleles datasets (n=5,554 pairs; **Fig. 1A**), second, paired TRA-HLA alleles (n=385 pairs; **Fig. 1A**) that contain TRA repertoires generated using Adaptive’s ImmunoSEQ assay, and third, TRA-HLA alleles dataset (n=855 pairs; **Fig. 1A**) containing repertoires profiled using MiLaboratories’ kits. For each of the three subsets we first split them randomly into 80% training and 20% validation sets to quantify the generalizable performance of the trained models on a comparable dataset. After that, we trained three classes of models on each of the three subsets and tested their performance on publicly available test datasets^13,14^ of paired TCR and HLA genotyping calls.

**Figure 1:**
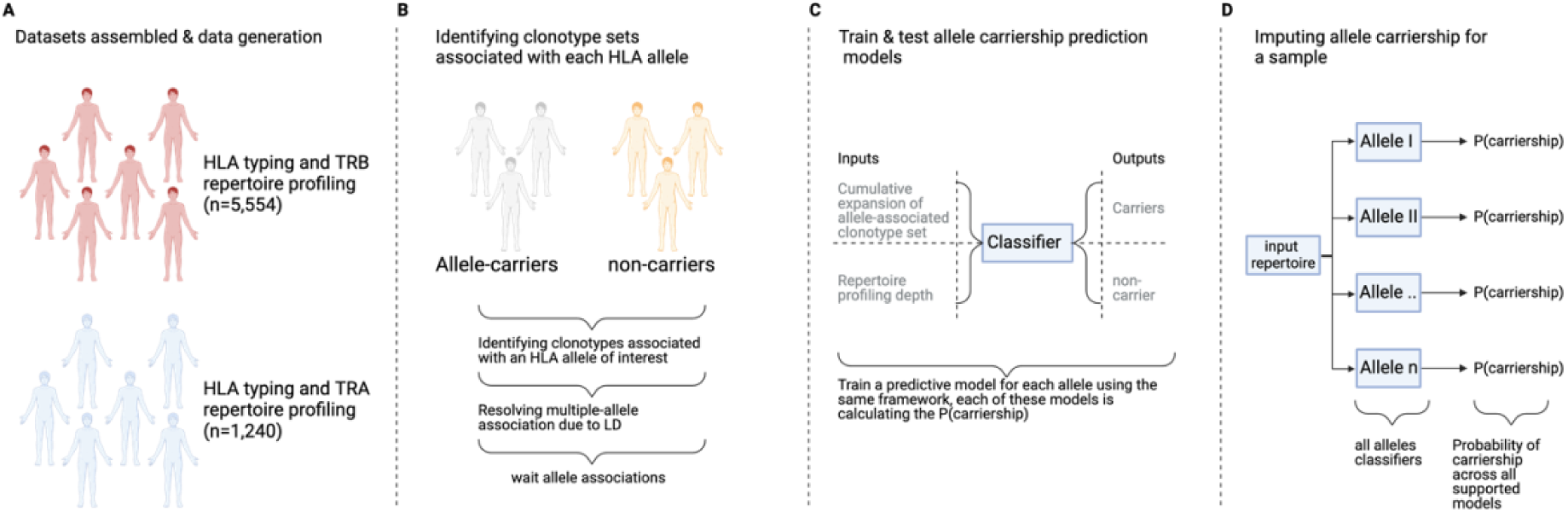
Overview of the approach used for discovering clonotypes restricted to different HLA proteins and for developing models to imputing the carriership of these alleles based on the TRA or the TRB repertoire. (**A**) shows the cohorts used in the current study to discover TRA- and TRB-clonotypes associated with different HLA alleles. (**B**) summarizes clonotype discovery associated with each allele by comparing the number of carriers and non-carriers using the Fisher’s exact test followed by resolving linkage-disequilibrium (LD) using L1LR models. (**C**) Summary of building a classifier to predict the carriership of alleles using the weight sum of the expansion of associated clonotypes of each allele and the profiled repertoire depth. (**D**) summarizes the pipeline for imputing HLA alleles from the provided repertoire, where for each of the supported allele models we calculate the carriership probability of each allele. The final HLA-calling represents alleles with a carriership probability of 0.5 or more.

**Figure 2:**
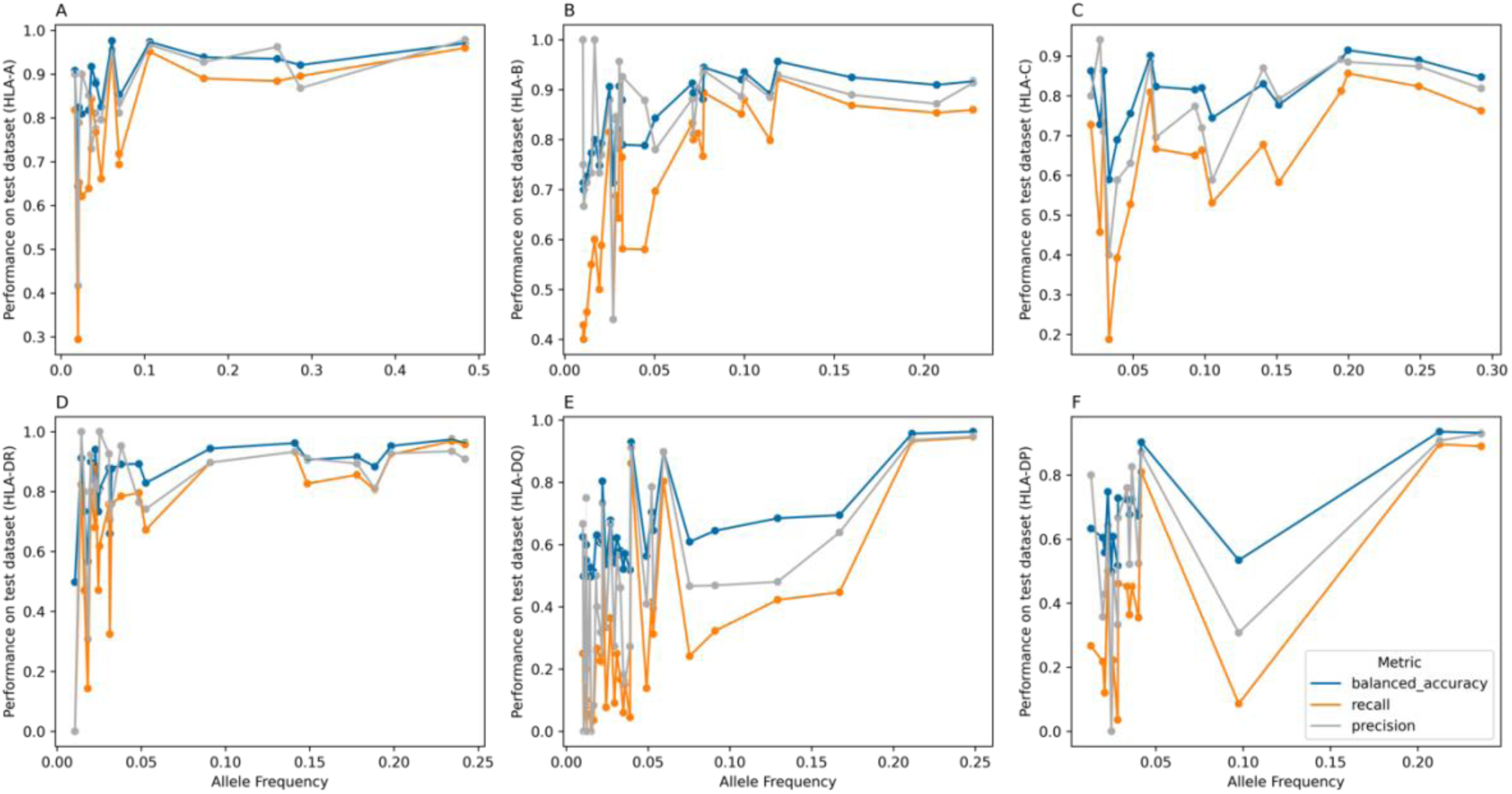
The relationship between HLA-allele carriership frequency and the performance of its TRB-based imputation model on a test dataset of 1,111 TRB repertoires with linked HLA calls. The performance metrics were used to evaluate the model performance, namely, balanced accuracy, recall and precision. (**A**-**C**) depict the performance of three HLA-I alleles, namely, HLA-A, HLA-B and HLA-C, respectively. Similarly, the performance of HLA-II alleles is illustrated in (**D**-**F**), with HLA-DR shown in (**D**), HLA-DQ in (**E**) and lastly, HLA-DP in (**F**).

**Table 1:** An overview of the datasets used for building HLA imputation models based on the T cell repertoire. HC refers to healthy controls, CD to individuals with Crohn’s disease and UC to individuals with ulcerative colitis. Lastly, IBD symptomatic controls represent individuals with symptoms of inflammatory bowel disease, but their endoscopy and lab tests failed to unambiguously diagnose CD or UC.

| DATASET NAME | LOCUS | NUMBER OF SAMPLES | COUNTRY | PHENOTYPE | TCR-SEQ METHOD |
| --- | --- | --- | --- | --- | --- |
| HC 1 | TRB | 773 | Germany | Healthy blood donors | Adaptive ImmunoSEQ |
| CD 1 | TRB | 1,186 | Germany | CD patients | Adaptive ImmunoSEQ |
| UC 1 | TRB | 480 | Germany | UC patients | Adaptive ImmunoSEQ |
| PSC 1 | TRB | 431 | Germany | PSC patients | Adaptive ImmunoSEQ |
| CD 2 | TRB | 1,809 | USA | CD patients | Adaptive ImmunoSEQ |
| UC 2 | TRB | 854 | USA | UC patients | Adaptive ImmunoSEQ |
| HC 2 | TRA | 165 | Germany | Healthy blood donors | Adaptive ImmunoSEQ |
| CD 3 | TRA | 124 | Germany | CD patients | Adaptive ImmunoSEQ |
| UC 3 | TRA | 96 | Germany | UC patients | Adaptive ImmunoSEQ |
| HC 3 | TRA | 264 | Norway | Symptomatic controls (IBD) | MiLaboratories kits |
| CD 4 | TRA | 231 | Norway | CD patients | MiLaboratories kits |
| UC 4 | TRA | 360 | Norway | UC patients | MiLaboratories kits |

### Discovering public TRB clonotypes associated with different HLA-alleles and developing HLA imputation models based on the TRB repertoire

After splitting the TRB-HLA dataset into 80% training and 20% validation we used a similar framework to Zahid *et al.*^11^ to build TRB-based HLA imputation models (**Online Methods; Fig. 1**). Briefly, we started by discovering clonotypes associated with HLA-alleles using the statistical framework described by Emerson *et al.*^9^. For each allele, the frequency of public clonotypes in allele carriers relative to non-carriers was compared using a one-sided Fisher’s exact test where clonotypes with an association P-value <1×10^-4^ are designated as clonotypes associated with the respective HLA allele (**Fig. 1B**). Subsequently, we used the L1-regularised linear regression model described by Zahid *et al.*^11^ to resolve the cases where the same clonotype is associated to multiple HLA alleles due to linkage-disequilibrium (**Fig. 1B)**. Then, we used a weighted sum of the abundance of allele-associated clonotypes as well as the repertoire depth to develop a linear regression model that can distinguish carriers of this allele from non-carriers^11^ (**Fig. 1C**). Lastly, by the running a given TRA or TRB repertoire across the models for supported HLA alleles, the HLA allele carriership of the sample can be inferred (**Fig. 1D**).

Using 80% of the data we were able to identify 722,060 clonotypes that were associated with 312 HLA alleles with the number of associated clonotypes being a function of allele frequency (**Fig. S1**). Subsequently, we restricted our analysis to alleles with carriership frequency >1%, leaving 600,095 unique clonotypes associated with 137 unique HLA alleles with a median of 2,211 clonotypes per HLA allele. Most of the clonotypes were restricted to HLA-II alleles (n=466,277), particularly, HLA-DRB1 (n=303,330) relative to all HLA-I alleles (n=145,224), potentially, because of the higher ratio of CD4^+^ T cells in the blood relative to CD8^+^ T cells. Using the L1-regularized logistic regression framework^11^ we were able to resolve the association between clonotypes and multiple HLA alleles, however, for a small subset of clonotypes this was not possible. Specifically, out of the 600,095 associated clonotypes, 587,224 (97.8%) clonotypes were associated with a single HLA allele while only 12,871 (2.2%) clonotypes were associated with multiple alleles.

After building prediction models for these 137 alleles, we tested their performance on the 20% validation dataset (n=1,111 paired TRB-HLA repertoire). Starting with HLA-A alleles, we observed a high performance across most alleles with a median balanced accuracy of 0.88, median precision of 0.851 and a median recall of 0.775 (**Fig. S2**). A similar trend was observed with HLA-B alleles where the median balanced accuracy was 0.87, and the median precision and recall were 0.88 and 0.76, respectively (**Fig. S3**). The performance of HLA-C allele models was lower than that of HLA-B and HLA-A, with a median balanced accuracy of 0.82, a median precision of 0.792, and a median recall of 0.66 (**Fig. S4**). This might be attributed to the small footprint of HLA-C on the TRB repertoire as it has a lower surface expression^15^ and a smaller immunopeptidome^16^.

Regarding HLA-II alleles, HLA-DR alleles illustrate a high performance relative to HLA-DQ and HLA-DP, with a median balanced accuracy of 0.89, median precision of 0.89 and a median recall of 0.79 (**Fig. S5**). HLA-DQ alleles had an average balanced accuracy of 0.60, an average precision of 0.46, and an average recall of 0.22 (**Fig. S6**). While the average model performance was generally inferior to other HLA proteins discussed so far, some HLA-DQ models showed higher performance. For example, HLA-DQA1*01:02-DQB1*06:02 showed a balanced accuracy of 0.96, precision of 0.94 and a recall of 0.94 (**Fig. S6**). A similar trend was also observed with HLA-DP alleles, where the average balanced accuracy was 0.67 and the average precision and recall were 0.58 and 0.35, respectively (**Fig. S7**). These performance metrics were mainly driven by a handful of alleles that showed an accurate predictive performance such as the HLA-DPA1*01:03-DPB1*04:02 model, which had a balanced accuracy of 0.93, a precision of 0.92 and a recall of 0.89 (**Fig. S7**).

To investigate factors influencing the performance of these models, we started by analyzing the impact of allele frequency and model performance. For HLA-A alleles we observed a positive correlation between the carriership frequency and different performance metrics (**Fig. 1A**). This relationship was not linear and showed signs of saturation, where carriership frequencies above 0.05-0.1 did not translate into a meaningful increase in the model performance. Similar findings were also observed for HLA-B (**Fig. 1B**), HLA-C (**Fig. 1C**) and HLA-DR (**Fig. 1D**) models. For HLA-DQ (**Fig. 1E**) and HLA-DP (**Fig. 1F**), this trend was also observed but at a weaker extent, with some alleles having relatively high carriership frequency (>0.1) but a relatively poor performance.

HLA-DQ and HLA-DP proteins are made from two different chains, α and β, leading to the formation of cis and trans proteins, with the cis proteins formed between the chains being encoded on the same chromosome while the trans complexes are formed between α and β chains encoded on different chromosomes, for example, an α chain encoded by the paternal copy and a β chain encoded by the maternal copy or *vice versa*. While the same applies to HLA-DR proteins, the α chain of the HLA-DR is invariant, and hence we focused only on the β chain encoded either by the paternal or the maternal chromosomal copy. Several studies have indicated that trans complexes have a minor impact on the formed immunopeptidome^17^, as not all trans complexes lead to a stable HLA protein formation at the cell surface^18^. This might explain the poor performance of different HLA-DQ and HLA-DP complexes that have relatively high carriership frequency, as these αβ allele combinations might not generate a stable HLA complex and hence have a minor impact on the TRB repertoire.

Motivated by these findings and the performance of the models on the validation dataset, we used the entire dataset for training and developing imputation models using the same workflow introduced above. This enabled us to identify 1,095,576 unique TRB clonotypes that were associated with 437 unique HLA alleles, with 1,049,766 clonotypes showing single-allele association and 45,810 being associated with multiple alleles. After filtering for alleles with carriership frequency >0.01 (corresponds to n>55 individuals), we obtained 891,564 clonotypes that were associated with 175 HLA alleles, specifically, with 17 HLA-A alleles, 27 HLA-B, 17 HLA-C, 22 HLA-DR, 30 HLA-DP, and 62 HLA-DQ alleles (**Table S1**). Similar to previous findings, the number of associated clonotypes was higher with HLA-II alleles, particularly HLA-DRB1 alleles relative to HLA-I alleles (**Table S1**). Despite the strong differences between the different HLA loci, within a single locus, allele carriership frequency strongly correlated with the number of identified TRB clonotypes as shown in **Fig. S8**.

To test the performance of the developed models we used a previously published dataset^13^ of 229 healthy and IBD individuals with paired HLA genotypes and immune repertoires, where we imputed the HLA of each sample using the TRB repertoire and compared the imputed results to the provided HLA alleles. Furthermore, we focused the analysis on models with an allele carriership in the training dataset above 0.05. Although this test dataset was generated with a different TCR-Seq method and from RNA instead of DNA, our developed TRB-models were able to impute common HLA alleles (**Fig. 3** & **Fig. S9**). Across the different loci we observed a significant reduction in the recall (**Fig. 3B** & **Fig. S9**) which might be attributed to differences in the TCR-Seq methodology used, as the models were trained on Adaptive ImmunoSEQ data and tested on repertoire profiled using MiLaboratories’ kits. The former assay generally enables a deeper repertoire profiling relative to the latter. Hence, this reduction in the profiling depth might explain the reduction in the ability of the models to recall alleles. Nonetheless, the precision of the models remained relatively unchanged (**Fig. 3C** & **Fig. S9)**.

**Figure 3:**
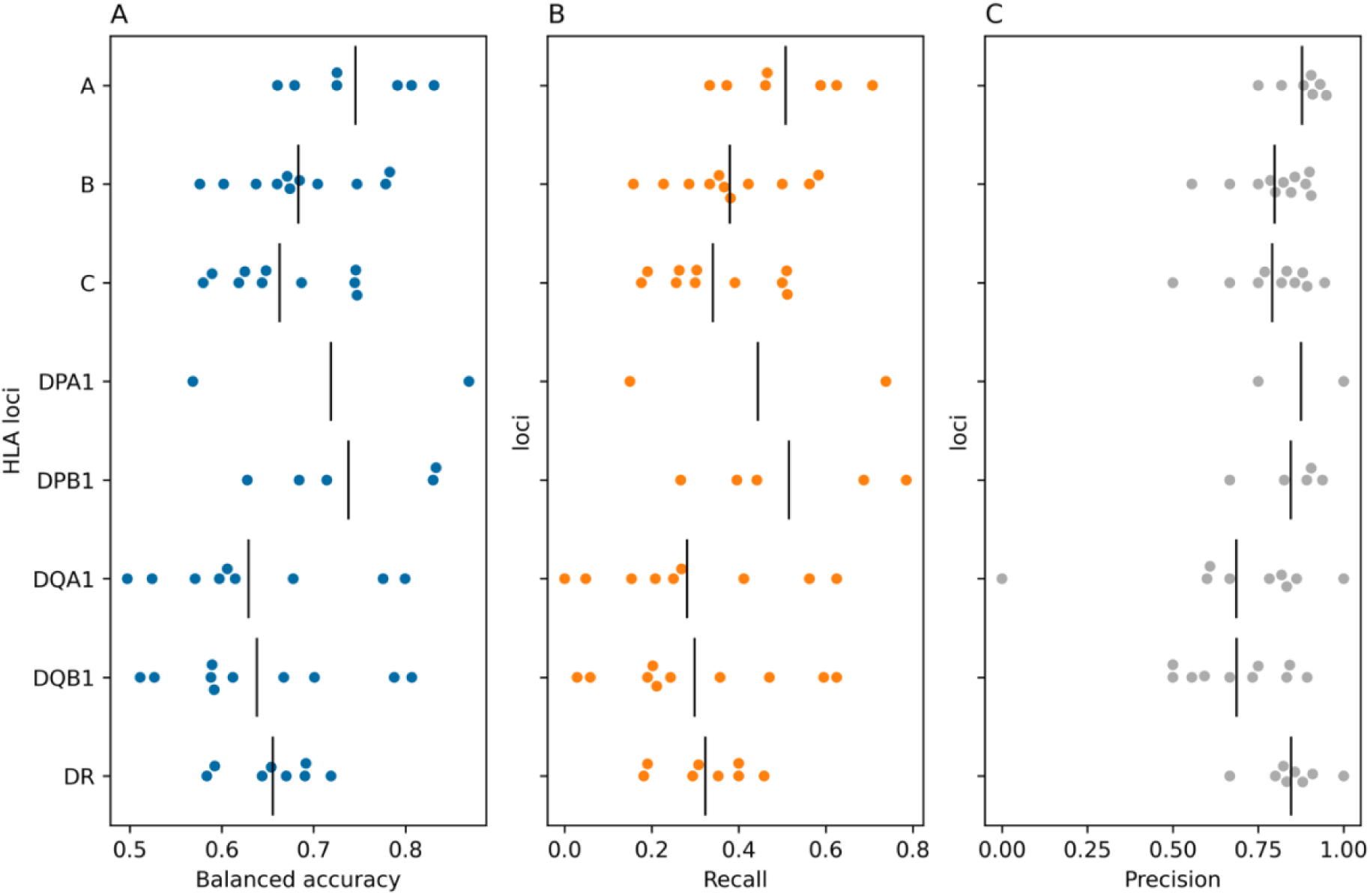
The performance of the TRB-based HLA imputation models on an independent test dataset obtained from Rosati et al.^13^. (**A**) shows the balanced accuracy, while (**B**) the recall and (**C**) the precision across different HLA alleles belonging the different HLA loci. Across all panels, alleles with carriership frequency <0.05 (n<12 samples) were excluded from the analysis.

To test the performance of the models on an independent test dataset that was generated with the same technology, *i.e.* Adaptive’s immunoSEQ, we used a subset of the immuneCODE^14^ database with matching HLA allele information (n= 63). Subsequently, we imputed the HLA allele of each sample from its TRB repertoire using the models and compare the results to the HLA typing results. As seen in **Fig. 4**, most of the models showed an accurate performance, except for some HLA-DQ complexes, which might be a trans HLA-DQ complex (**Fig. S10**). Also, by utilizing the same TCR-Seq technology, we observed a robust increase in the recall relative to the Rosati *et al.*^13^ test dataset. Indicating that the decrease in the recall observed previously can be attributed to the shallow profiling of the repertoire performed in the Rosati *et al.*^13^ study.

**Figure 4:**
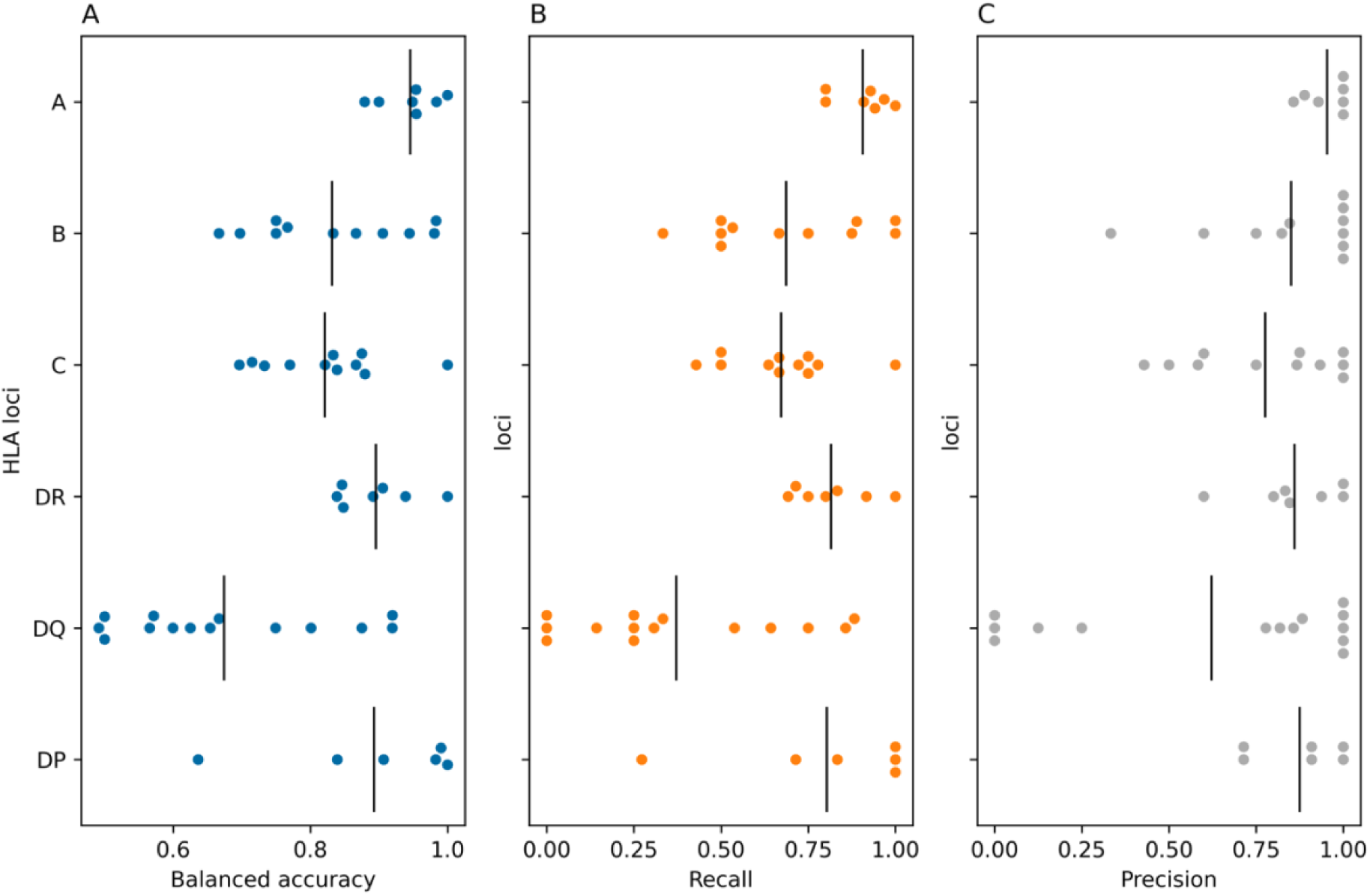
The performance of the TRB-based HLA imputation models on an independent test dataset obtained from the immuneCODE dataset^14^. (**A**) shows the balanced accuracy, while (**B**) the recall and (**C**) the precision across different HLA alleles belonging the different HLA loci. Across all panels, alleles with carriership frequency <0.05 (n<3 samples) were excluded from the analysis.

### Discovering public TRA clonotypes associated with different HLA-alleles and developing HLA imputation models based on the TRA repertoire

As the TRA repertoire was profiled using two different TCR-Seq technologies, namely Adaptive’s ImmunoSEQ and with MiLaboratories’ kits and using different starting materials, *i.e.* DNA and RNA, respectively, we did not combine the two datasets and treated each dataset independently. Starting with the dataset made of cohorts HC2, CD3 and UC3 which were profiled using Adaptive’s ImmunoSEQ from DNA (**Online Methods)**, we split the dataset into an 80% training and a 20% validation sets. We then used the framework described above to discover TRA clonotypes associated with HLA proteins (**Online Methods**). Although our results mirrored the results identified from the TRB, in which the number of associated clonotypes per allele was strongly dependent on the HLA allele frequency (**Fig. S11**), the overall number of TRA-associated clonotypes was lower than the number of TRB-associated clonotypes. This can be attributed to differences in size between the two datasets (>5,554 TRB repertoires vs. 308 TRA repertoires used here for training, *i.e.* ∼6% of the TRB dataset) which highlights the impact of the dataset size on discovering clonotypes associated with each allele. Given the small number of repertoires and discovered allele associated-clonotypes, the resulting TRA-based imputation models showed a relatively poor predictive performance, relative to the TRB-models, even for the very common alleles, *i.e.* alleles with carriership frequency >5% (**Fig. S12-S17**).

We repeated the same process with the other TRA-HLA datasets composite of HC3, CD4 and UC4 cohorts which were profiled using MiLaboratories’s kits with RNA as a starting biological material (**Online Methods**). Given that this dataset is ∼two-fold the size of the previous TRA-HLA dataset we observed a higher number of HLA-associated clonotypes, 31,230 relative to 9,435 clonotypes. Similar to previous findings, the number of clonotypes associated with each allele positivity correlated with allele frequency and was different among the different HLA loci (**Fig. S18**). After testing the models on the 20% validation dataset, we observed a similar trend to the TRB findings where HLA-A (**Fig. S19**) and HLA-B (**Fig. S20**), showed, on average, a higher performance relative to HLA-C (**Fig. S21**) across common HLA alleles (carriership frequency >0.05). For common HLA-II alleles, HLA-DR alleles showed the highest performance (**Fig. S22**) relative to HLA-DQ (**Fig. S23**) and HLA-DP (**Fig. S24**). Although the performance of these models was relatively higher than the first TRA-based models trained on the smaller TRA dataset obtained using immunoSEQ, it is still inferior to the TRB-based models in terms of the number of supported HLA-alleles and the accuracy of each model.

Consequently, we focused the TRA-based model development on the larger dataset assembled from HC3, CD4 and UC4 cohorts (n=855 TRA-HLA pairs) where we used the entire dataset to develop imputation models for HLA alleles with a carriership frequency above 0.05. To test the generalizability of these models we used the independent Rosati *et al.*^13^ test dataset (**Fig. 5 & Fig. S25**). Relative to TRB-based imputation models, the TRA-models showed an inferior predictive performance potentially due to the much smaller training dataset, 855 TRA-HLA, relative to 5,554 TRB-HLA pairs. Similar to TRB-based models, the TRA-based models suffered from a reduction in recall relative to precision, potentially due to the shallow repertoire depth of the Rosati *et al.*^13^ dataset.

**Figure 5:**
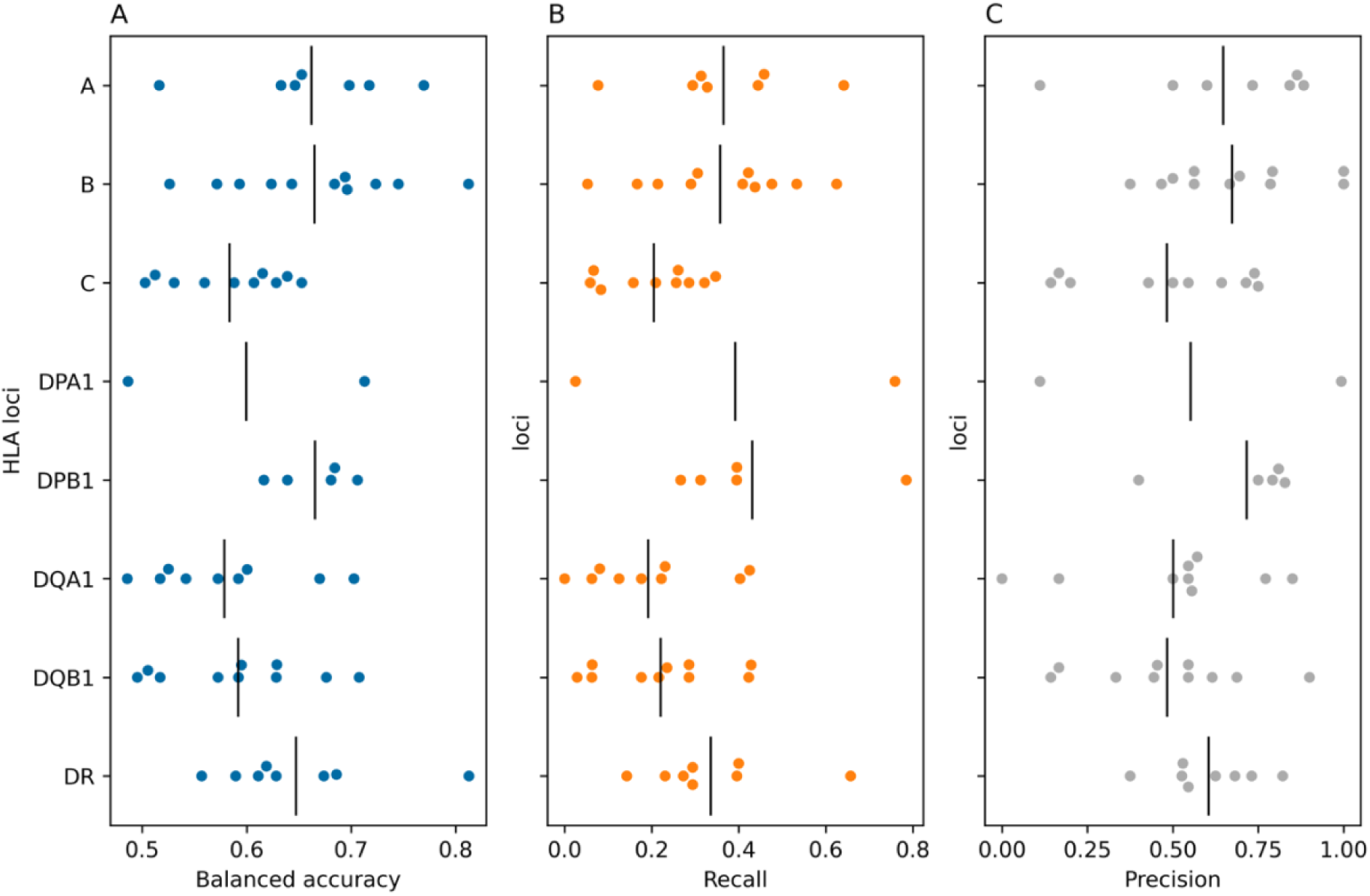
The performance of the developed TRA-based imputation models on a test dataset of paired TRA repertoire and HLA alleles that was generated by Rosati et al.^13.^ (**A**) shows the balanced accuracy, while (**B**) the recall and (**C**) the precision across different HLA alleles belonging the different HLA loci. Across all panels, alleles with carriership frequency <0.05 (n<12 samples) were excluded from the analysis.

### The discovered HLA-associated clonotypes target common infections

To gain more insights into the identified TRA- and TRB-clonotypes that are associated with different HLAs, we analyzed their overlap with public TCR-antigen databases, namely, McPAS^19^ and VDJdb^20^. Out of the 54,300 TRB clonotypes defined in these databases, 1,910 clonotypes overlapped between the two datasets, most of the identified clonotypes were targeting common infections such as cytomegalovirus (CMV), Epstein-Barr virus (EBV) and influenza. Furthermore, most of the identified clonotypes were restricted to common HLA-I alleles such as HLA-A*02:01, HLA-A*03:01, HLA-B*08:01 and HLA-B*07:02. Utilizing a sequence clustering analysis on the overlapped clonotypes (**Online Methods**), we did not observe a significant degree of sequence similarity among the identified clonotypes (**Fig. 6A**), suggesting that the identified clonotypes are recognizing different antigens presented by the same HLA protein.

**Figure 6:**
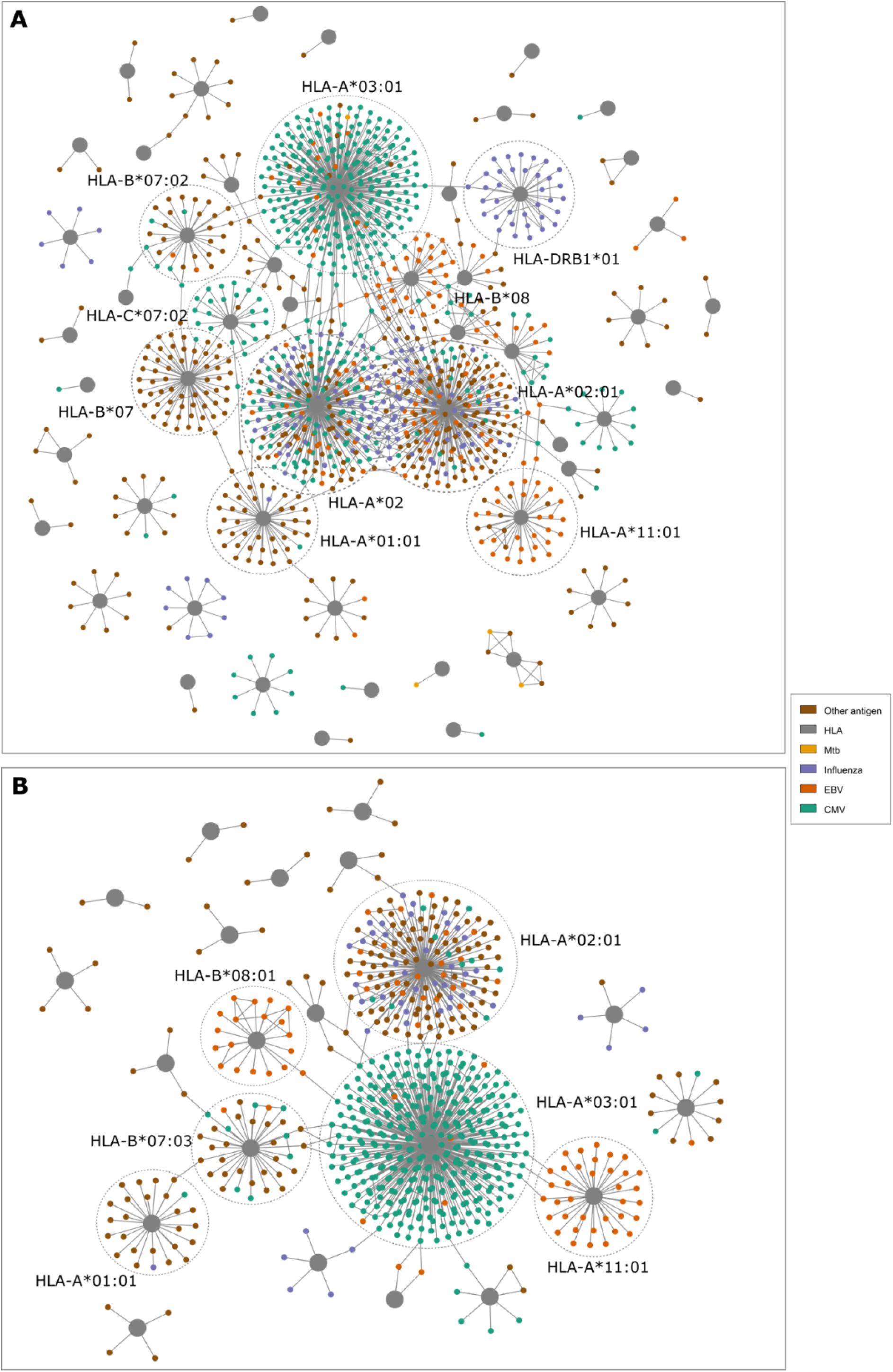
Antigenic specificity of HLA associated TRA- and TRB-clonotypes. (**A**) depicts the overlap between HLA-associated TRB clonotypes and public databases, namely, VDJdb^20^ and McPAS^19^ while (**B**) illustrates the overlap between these databases and HLA-associated TRA clonotypes. In (**A**) and (**B**), clusters with multiple clonotypes are annotated with the associated HLA allele. Network visualization was performed using Cytoscape^21^.

Afterward, we investigated the identified TRA clonotypes that are HLA associated using the same dataset described above^19,20^. Out of the 55,293 clonotypes identified from both TRA-HLA datasets, 1,049 clonotypes overlapped with the assembled public database (n= 26,937). Similar to the findings of TRB clonotypes, most of the identified clonotypes were restricted to common viral and bacterial infections such as CMV, EBV, influenza, and *M. tuberculosis*. Most of the overlapped clonotypes were specific toward HLA-I restricted antigens (904; 86%) with only 16 clonotypes (1.5%) being specific to HLA-II alleles. Across these annotated peptides the majority were restricted to common HLA-A alleles, such as HLA-A*02:01 and HLA-A*03:01 proteins. Mirroring the TRB-based analysis, we did not observe a significant degree of sequence similarity among the identified clonotypes within the annotated clusters of each HLA protein (**Fig. 6B**).

## Discussion

We assembled a large dataset of paired TCR repertoires with their corresponding HLAs which enabled us to discover thousands of clonotypes that are associated with hundreds of HLA alleles. Based on these clonotypes we were able to develop robust statistical models for imputing common HLA alleles. While previous studies focused mainly on analyzing the relationship between the TRB repertoire and HLA proteins^9,11^, using 1,240 TRA-HLA pairs, we illustrate that the TRA repertoire can also be used to impute HLA alleles from the TRA repertoire. Although the assembled TRA-HLA dataset reported here represents, to the best of our knowledge, the biggest dataset of its kind, it still represents only (<25%) of the TRB-HLA dataset. This also explains the inferior performance of TRA-based relative to TRB-based imputation models, highlighting the critical impact of dataset size on the predictive performance of these imputation models and generally on studying the interplay between TCRs and HLA proteins.

Besides the size of the dataset, the utilized TCR-Seq method impacts the imputation results. TCR-Seq can be conducted using different methods such multiplex PCR, and 5’-RACE, using DNA or RNA, and using different quantities of the provided genetic materials^6^, all of which have an impact on shaping the identified T cell repertoire^22^. The performance of the developed TRA and TRB imputation models depends on the utilized TCR-Seq as the models showed superior performance with the immuneCODE test dataset^14^ relative to the test dataset reported by Rosati *et al.*^13^ which was generated using a different TCR-Seq method. Lastly, both of our TRA- and TRB-based imputation models are based on the blood repertoire. We have recently shown systematic differences in the composition of the blood repertoire relative to tissues^23^. Thus, the performance of these models on the immune repertoire of tissues would vary based on their similarity to the blood repertoire for which they were developed.

Despite the large size of the TRB-HLA dataset utilized in the current study, we were able to develop accurate models for HLA proteins that are common among European and Northen American populations only. This is because the performance of each of these models is strongly dependent on the corresponding HLA allele frequency in the training dataset. The relationship between allele frequency and different performance metrics followed a dose-response curve with a marginal improvement in performance once the allele carriership frequency was >0.1. Thus, building models, or even discovering associated clonotypes, for rare HLA alleles was not possible. These rare HLA alleles might exhibit a higher frequency in different populations, for example, HLA-DRB1*15:02 and HLA-DRB1*15:03 alleles are rarer in European-populations but higher in Asian and sub-Saharan African populations^24^. Selective sampling can be used to mitigate this problem, where the TCR repertoire is preferentially profiled for individuals with rare HLA-alleles, to provide a cost-effective way to extend the number of HLA alleles supported.

Another potential limitation to our study is that most of the individuals included in the study were suffering from different forms of inflammatory diseases, mainly, IBD. Our dataset is therefore enriched for known disease-associated HLA alleles such as HLA-DRB1*15:01^24^ and HLA-DRB1*07:01^25^, however, these alleles are also common in healthy individuals. A second limitation is using genotyping-based HLA-imputation to generate paired TCR-HLA database for training and for developing TRA- and TRB-based HLA imputation models. Given that we focused only on common alleles, for which the genotyping-based imputation is fairly accurate^12^, the noise introduced in the TCR-HLA association database should be minimal.

While HLA allele frequency was an important factor in determining the number of clonotypes associated with each HLA allele, there was a notable difference among the HLA proteins encoded by different loci. HLA-A, HLA-B and HLA-DR alleles had on average a higher number of associated clonotypes relative to HLA-C, HLA-DQ and HLA-DP alleles. Different factors can contribute to this such as the level of surface expression where HLA-C has been shown to have a lower surface expression relative to other HLA-I proteins^15^. Similarly, HLA-DP and HLA-DQ have a lower surface expression relative to HLA-DR proteins^26^. Beyond the surface expression, the complexity and diversity of the immunopeptidome of these different HLA alleles have been shown to vary considerably, for example, the complexity of the HLA-C immunopeptidome is lower than for HLA-A and HLA-B immunopeptidomes^16^. These findings indicate, at least from the T cell perspective, that antigen presentation by HLA-A, HLA-B and HLA-DR plays an important role in shaping adaptive immune responses relative to other HLA proteins.

Although we could not resolve the antigenic specificity for all HLA-associated clonotypes using public databases^19,20^, we were able to annotate the antigenic specificity for >1,000 TRA and TRB clonotypes. Most of these clonotypes were specific to common viral and bacterial infections. Thus, the identified allele-associated clonotypes represent a fraction of the immune memory toward prevalent infections. Mechanistically, the interaction between infections and HLA proteins will lead to the formation of different HLA-peptide complexes in different individuals, these complexes will be recognized by different T cells. Hence, carriers of a given allele will have a shared response in terms of T cell clonotypes toward the same HLA-peptide complex relative to individuals with different HLA alleles. Thus, the subset of allele-associated clonotypes present in an individual represent a fraction of the immune exposure history of this individual. By decoding the antigenic specificity of these clonotypes a better understanding of the immunological history of an individual and/or a population can be developed. Alterations in responses to some of these common infections have been associated with different diseases, for example, inappropriate immune responses toward Epstein-Barr virus have been associated with multiple autoimmune and chronic inflammatory diseases such as multiple sclerosis^27^, rheumatoid arthritis^28^, and Sjögren syndrome^29^. Additionally, cytomegalovirus has been linked with the adult-onset Still’s disease^30^, and human papillomaviruses with cervical cancers^31^. It should be mentioned that the public databases such as McPAS^19^ and VDJdb^20^ are far from complete and are biased towards, common viral pathogens. Therefore, more systematic efforts that decode the antigenic specificity of TCRs are needed to enrich the databases with the antigenic specificity of public T cell clonotypes. Different techniques can be used to investigate the antigenic specificities of these clonotypes such as yeast-display^32^, phage-display^33^ and receptor–antigen pairing by targeted retroviruses (RAPTR)^34^, among others.

In conclusion, our results indicate the utility of coupling statistical learning with paired population-level profiling of TCR repertoire and HLA genotypes in terms of identifying sets of public clonotypes that are restricted to common HLA alleles. Having the presenting HLA candidates at hand for a set of TCR of interest (*e.g.* because these TCRs were identified as disease-specific) opens new experimental avenues. Although we illustrate here that these clonotypes can be used to infer the HLA background of a sample, they are also marker of previous common infections. By decoding their antigenic specificity, a better understanding of responses to these infectious agents at both the individual and population level can be elucidated.

## Acknowledgments

We would also like to thank the Crohn’s & Colitis Foundation IBD Plexus program for providing us with the T cell repertoire profiles and the genotypes of the SPARC IBD cohort. We would like to also thank Sören Franzenburg, Janina Fuß, Rebekka Kraemer, Nicole Braun, Maria Eloina Figuera Basso, Anja Tanck, Yewgenia Dolshanskaya, and Melanie Vollstedt for their help with T cell repertoire profiling. We would also like to thank Michel Wittig and Tanja Wesse for their help with SNP array genotyping and for Christoph Prieß and Lars Wienbrandt for providing computational support to the project.

## Online Methods

### Ethical approval and sample collection

The study has been approved by the ethical committee at the University of Kiel under the following ethical votes: D441/16, D474/12, A161/08, A103/14, and A148/14. For the Norwegian cohort, namely, H3, CD4, and UC4 datasets were derived from the IBSEN III study which was approved by the South-Eastern Regional Committee for Medical and Health Research Ethics (Ref 2015/946-3) and performed in accordance with the Declaration of Helsinki. The USA-based samples, namely, UC2 and CD2 are derived from the SPARC IBD cohort from the IBD Plexus research program maintained by the Crohn’s & Colitis Foundation and described by Raffals *et al.*^35^ A written informed consent was collected from all participants prior to the beginning of the study.

### T cell repertoire profiling

Two technologies were used for profiling the T cell receptor, namely, Adaptive ImmunoSEQ and MiLaboratories kits. Adaptive ImmunoSEQ (Adaptive Biotechnologies) assays were conducted as described previously^36,37^ using up to 18 ug of peripheral blood DNA. While MiLaboratories kits were conducted according to the manufacturer’s instructions. Briefly, 300ng of RNA per samples isolated from PAXgene tubes were used to profile the T cell repertoire of peripheral blood. After library preparation, sequencing was conducted on a NovaSeq X with an average of 5 million reads per chain per sample. Lastly, clonotypes were assembled from the sequencing data using MIXCR^38^. For both TCR-Seq technologies, we started processing the samples by filtering non-productive clonotypes, *e.g.* clonotypes that contain a stop codon, or a frameshift mutation. After that, different V(D)J rearrangements with the same V and J genes as well as CDR3 amino acid sequences were collapsed into a single clonotype. Hence, the number of clonotypes is synonymous with the number of unique functional V(D)J recombination events.

### Genotyping and HLA imputation

All samples were genotyped using Illumina Infinium Global Screening Array beadchips (GSAMD-24v1-0_20011747_A1 or GSAMD-24v2-0_20024620_A1). Genotype quality control (QC) was conducted independently for each cohort, following the procedures outlined in BIGWAS quality control pipeline^39^. In short, the pipeline performs chip type detection and normalizes variant names. Variants with a missingness rate exceeding 0.02 were excluded from further analysis. Additionally, variants are filtered by Hardy-Weinberg equilibrium with a threshold of p=10^-5^. Sample QC was disabled in BIGWAS QC with the parameter ‘--skipsampleqc=1’. To facilitate population structure analysis, each cohort was combined with 2,504 samples from the 1000 Genomes Project for principal component analysis (PCA). Samples were projected onto the first two principal components, and those falling within the median ± 5 times the interquartile range (IQR) along these components were identified as Europeans.

HLA imputation was performed with the HLApipePublic pipeline available at https://github.com/ikmb/HLApipePublic. In brief, the pipeline filters the input dataset to variants within the region of 29mb and 34mb of chromosome 6 and aligns them to the imputation reference. Imputation of HLA alleles was calculated using the HIBAG algorithm^40^, which applies attribute bagging (BAGging) to enhance prediction accuracy. The analysis utilized the multi-ethnic IKMB reference panel, as described in^12^. Phasing of alleles is done with the tool SHAPEIT2^41^. This approach generates predictions by averaging HLA-type posterior probabilities across an ensemble of classifiers built on bootstrap samples^42^. Imputation was performed at full four-digit resolution for the following loci: HLA-A, HLA-C, HLA-B, HLA-DRB3, HLA-DRB5, HLA-DRB4, HLA-DRB1, HLA-DQA1, HLA-DQB1, HLA-DPA1, and HLA-DPB1.

### Identifying HLA-allele associated clonotypes

We followed the same framework developed by Emerson *et al.*^9^ to identify disease associated clonotypes. For each allele in each of the six classical HLA proteins, namely, HLA-A, HLA-B, HLA-C, HLA-DR, HLA-DQ and HLA-DP we binned samples into two categories, carriers and non-carriers. Subsequently, we compared the incidence of each public clonotypes, defined as clonotypes detected in two individuals or more, in carriers and non-carriers. Thus, for each clonotype we end up with a 2×2 contingency table that contains the number of individuals that have this allele and this clonotypes, have this clonotypes but not the allele, have the allele but not the clonotypes, or does not have neither the allele nor the clonotypes. After building the table, we used a one-sided Fisher exact test to investigate the statistical association between the clonotypes and the HLA alleles. Although Zahid *et al.*^11^ optimized the significance cutoff for different allele, due to the computationally intensive nature of this approach we used a fixed cutoff of 1×10^-4^ to define clonotypes that are associated with a particular allele.

### Developing the TCR-based HLA imputation framework

We used a similar approach to Zahid *et al.*^11^ were we started by resolving the promiscuous association problem where a single clonotypes is associated to multiple HLA alleles. Although we observed this problem among alleles of the same gene, it was much more apparent among alleles of different genes. A potential cause for this problem is linkage disequilibrium (LD) where due to the statistical association nature of the study, we identify clonotypes that are associated with a particular haplotype instead of a particular HLA allele. To resolve this, we followed the L1 regularized linear regression framework proposed by Zahid *et al.*^11^. However, instead of building models for each clonotype using the HLA alleles with a strong LD as done by Zahid *et al.*^11^ we included all the HLA alleles associated with a clonotype. Subsequently, we arrange the dataset into HLA-alleles and their corresponding HLA associated clonotypes. For each clonotype in the set we can calculated a weight that can be interpret as the strength of its association with its cognate allele following the same recipe described by Zahid *et al.*^11^. These weights to calculate a weighted sum of the expansion of allele-associated clonotypes in each repertoire, Lastly, used a logistic regression classifier to classify repertoire into allele carrier based on the number of unique clonotypes and the weighted sum of the expansion of allele-associated clonotypes in this samples as described by Zahid *et al.*^11^.

### Clustering of HLA-associated clonotypes

In order to cluster HLA-associated clonotypes, we followed a graph-based approach in which each clonotype is defined as a node. To draw an edge between these nodes, these two nodes need to have the same V and J genes, in addition to having at most, 1-hamming distance between the CDR3 of their amino acid sequences.

## References

1. Raskov, H., Orhan, A., Christensen, J. P. & Gögenur, I. Cytotoxic CD8+ T cells in cancer and cancer immunotherapy. Br J Cancer 124, 359–367 (2021).

2. Weisberg, S. P. et al. Tissue-Resident Memory T Cells Mediate Immune Homeostasis in the Human Pancreas through the PD-1/PD-L1 Pathway. Cell Rep 29, 3916–3932.e5 (2019).

3. Kjer-Nielsen, L. et al. MR1 presents microbial vitamin B metabolites to MAIT cells. Nature 491, 717–723 (2012).

4. Beckman, E. M. et al. Recognition of a lipid antigen by CD1-restricted αβ+ T cells. Nature 372, 691–694 (1994).

5. Laplagne, C. et al. Self-activation of Vγ9Vδ2 T cells by exogenous phosphoantigens involves TCR and butyrophilins. Cell Mol Immunol 18, 1861– 1870 (2021).

6. Mahdy, A. K. H. et al. Bulk T cell repertoire sequencing (TCR-Seq) is a powerful technology for understanding inflammation-mediated diseases. J Autoimmun 149, 103337 (2024).

7. Sharon, E. et al. Genetic variation in MHC proteins is associated with T cell receptor expression biases. Nat Genet 48, 995–1002 (2016).

8. Ishigaki, K. et al. HLA autoimmune risk alleles restrict the hypervariable region of T cell receptors. Nat Genet 54, 393–402 (2022).

9. Emerson, R. O. et al. Immunosequencing identifies signatures of cytomegalovirus exposure history and HLA-mediated effects on the T cell repertoire. Nat Genet 49, 659–665 (2017).

10. Ortega, M. R. et al. Learning predictive signatures of HLA type from T-cell repertoires. bioRxiv 2024.01.25.577228 (2024) doi:10.1101/2024.01.25.577228.

11. Zahid, H. J. et al. Large-scale statistical mapping of T-cell receptor β; sequences to Human Leukocyte Antigens. bioRxiv 2024.04.01.587617 (2024) doi:10.1101/2024.04.01.587617.

12. Degenhardt, F. et al. Construction and benchmarking of a multi-ethnic reference panel for the imputation of HLA class I and II alleles. Hum Mol Genet 28, (2019).

13. Rosati, E. et al. A novel unconventional T cell population enriched in Crohn’s disease. Gut 71, 2194 LP – 2204 (2022).

14. Nolan, S., et al. A large-scale database of T-cell receptor beta (TCRβ) sequences and binding associations from natural and synthetic exposure to SARS-CoV-2. Res Sq (2020).

15. Vince, N. et al. HLA-C Level Is Regulated by a Polymorphic Oct1 Binding Site in the HLA-C Promoter Region. The American Journal of Human Genetics 99, 1353– 1358 (2016).

16. Schellens, I. M. M. et al. Comprehensive Analysis of the Naturally Processed Peptide Repertoire: Differences between HLA-A and B in the Immunopeptidome. PLoS One 10, e0136417- (2015).

17. Nilsson, J. B. et al. Machine learning reveals limited contribution of trans-only encoded variants to the HLA-DQ immunopeptidome. Commun Biol 6, 442 (2023).

18. Kwok, W. W., Kovats, S., Thurtle, P. & Nepom, G. T. HLA-DQ allelic polymorphisms constrain patterns of class II heterodimer formation. J Immunol 150, 2263–2272 (1993).

19. Tickotsky, N., Sagiv, T., Prilusky, J., Shifrut, E. & Friedman, N. McPAS-TCR: a manually curated catalogue of pathology-associated T cell receptor sequences. Bioinformatics 33, 2924–2929 (2017).

20. Goncharov, M. et al. VDJdb in the pandemic era: a compendium of T cell receptors specific for SARS-CoV-2. Nat Methods 19, 1017–1019 (2022).

21. Shannon, P. et al. Cytoscape: a software environment for integrated models of biomolecular interaction networks. Genome Res 13, 2498–2504 (2003).

22. Barennes, P. et al. Benchmarking of T cell receptor repertoire profiling methods reveals large systematic biases. Nat Biotechnol 39, 236–245 (2021).

23. Mahdy, A. K. H. et al. Simultaneous Profiling of the Blood and Gut T and B Cell Repertoires in Crohn’s Disease and Symptomatic Controls Illustrates Tissue-specific Alterations in the Immune Repertoire of Crohn’s Disease Patients. bioRxiv 2024.09.22.614337 (2024) doi:10.1101/2024.09.22.614337.

24. Degenhardt, F. et al. Trans-ethnic analysis of the human leukocyte antigen region for ulcerative colitis reveals shared but also ethnicity-specific disease associations. Hum Mol Genet (2021) doi:10.1093/hmg/ddab017.

25. El Hadad, J., Schreiner, P., Vavricka, S. R. & Greuter, T. The Genetics of Inflammatory Bowel Disease. Mol Diagn Ther 28, 27–35 (2024).

26. Grifoni, A. et al. Characterization of Magnitude and Antigen Specificity of HLA-DP, DQ, and DRB3/4/5 Restricted DENV-Specific CD4+ T Cell Responses. Front Immunol 10, (2019).

27. Bjornevik, K. et al. Longitudinal analysis reveals high prevalence of Epstein-Barr virus associated with multiple sclerosis. Science (1979) 375, 296–301 (2022).

28. Fechtner, S. et al. Antibody Responses to Epstein-Barr Virus in the Preclinical Period of Rheumatoid Arthritis Suggest the Presence of Increased Viral Reactivation Cycles. Arthritis & Rheumatology 74, 597–603 (2022).

29. Pasoto, S. G. et al. EBV reactivation serological profile in primary Sjögren’s syndrome: an underlying trigger of active articular involvement? Rheumatol Int 33, 1149–1157 (2013).

30. Jia, J. et al. Cytomegalovirus Infection May Trigger Adult-Onset Still’s Disease Onset or Relapses. Front Immunol 10, (2019).

31. Okunade, K. S. Human papillomavirus and cervical cancer. J Obstet Gynaecol (Lahore) 40, 602–608 (2020).

32. Wang, L. & Lan, X. Rapid screening of TCR-pMHC interactions by the YAMTAD system. Cell Discov 8, 30 (2022).

33. Ch’ng, A. C. W., Lam, P., Alassiri, M. & Lim, T. S. Application of phage display for T-cell receptor discovery. Biotechnol Adv 54, 107870 (2022).

34. Dobson, C. S. et al. Antigen identification and high-throughput interaction mapping by reprogramming viral entry. Nat Methods 19, 449–460 (2022).

35. Raffals, L. E. et al. The Development and Initial Findings of A Study of a Prospective Adult Research Cohort with Inflammatory Bowel Disease (SPARC IBD). Inflamm Bowel Dis 28, 192–199 (2022).

36. Carlson, C. S. et al. Using synthetic templates to design an unbiased multiplex PCR assay. Nat Commun 4, 2680 (2013).

37. Gittelman, R. M., et al. Longitudinal analysis of T cell receptor repertoires reveals shared patterns of antigen-specific response to SARS-CoV-2 infection. JCI Insight 7, (2022).

38. Bolotin, D. A. et al. MiXCR: software for comprehensive adaptive immunity profiling. Nat Methods 12, 380–381 (2015).

39. Kässens, J. C., Wienbrandt, L. & Ellinghaus, D. BIGwas: Single-command quality control and association testing for multi-cohort and biobank-scale GWAS/PheWAS data. Gigascience 10, giab047 (2021).

40. Zheng, X. et al. HIBAG—HLA genotype imputation with attribute bagging. Pharmacogenomics J 14, 192–200 (2014).

41. Delaneau, O., Zagury, J.-F. & Marchini, J. Improved whole-chromosome phasing for disease and population genetic studies. Nat Methods 10, 5–6 (2013).

42. Zheng, X. et al. HIBAG-HLA genotype imputation with attribute bagging. Pharmacogenomics J. 14, 192–200 (2014).

