## Supplementary figures & Tables for "Decoding the restriction of T cell receptors to human leukocyte antigen alleles using statistical learning"

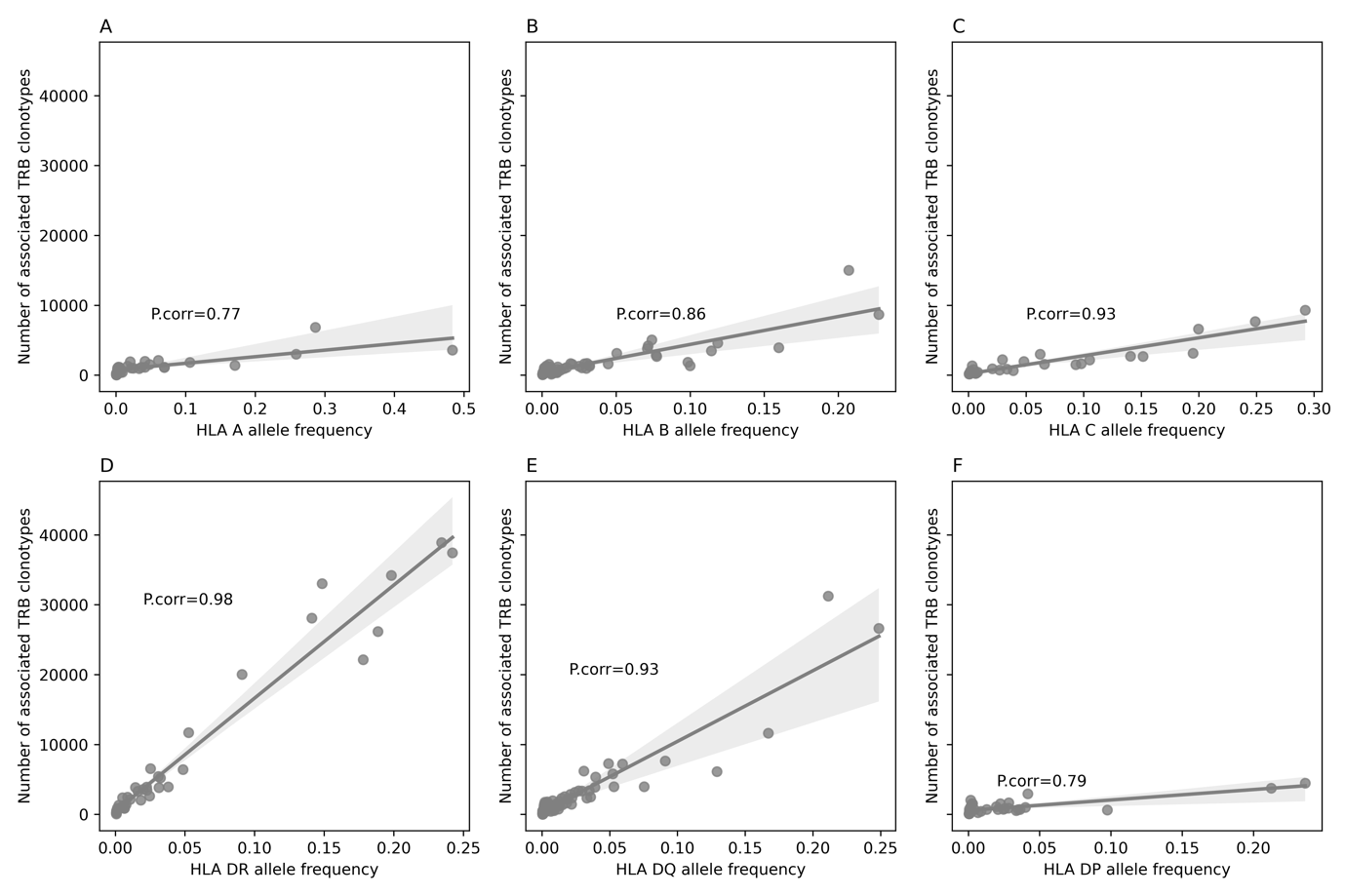

**Figure S1**: The relationship between HLA allele carriership frequency (shown on the x-axis) and the number of associated TRB clonotypes (shown on the y-axis) for the six classical HLA proteins, namely, HLA-A (**A**), HLA-B (**B**), HLA-C (**C**), HLA-DR (**D**), HLA-DQ (**E**), HLA-DP (**F**). Lastly, P.corr donates the Pearson correlation co-efficiency.

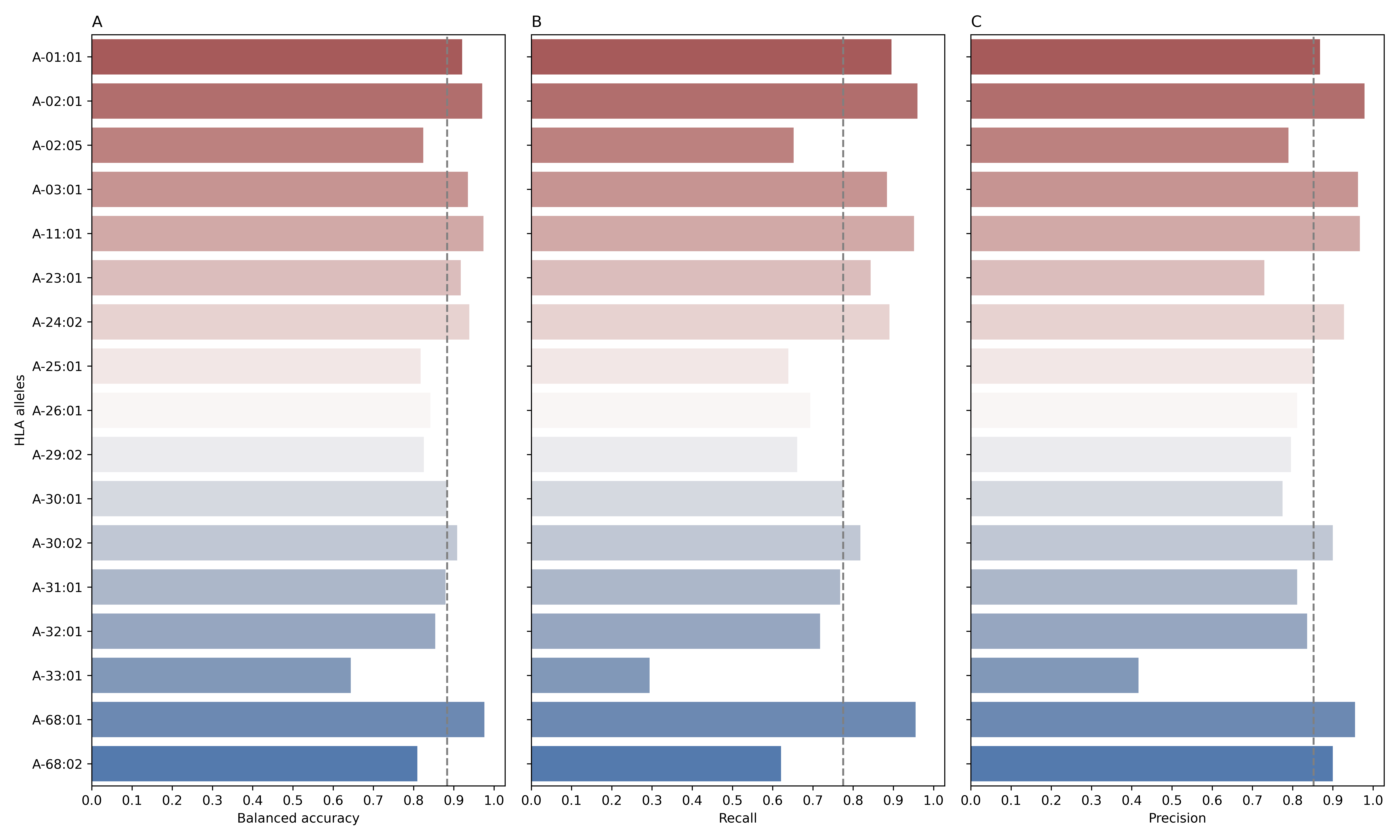

**Figure S2**: The performance of different models to predict the HLA-A allele-carriership status from the TRB repertoire evaluated on a test dataset made of 1,111 repertoires. On the y-axis the models of each HLA-A allele are shown, for example, A-01:01 represents a model that predicts whether an individual is a carrier for the HLA-A*01:01 allele or not. (**A**), (**B**) and (**C**) depict the balanced accuracy, recall and precision, respectively. Across all panels, grey dashed lines represent the median.

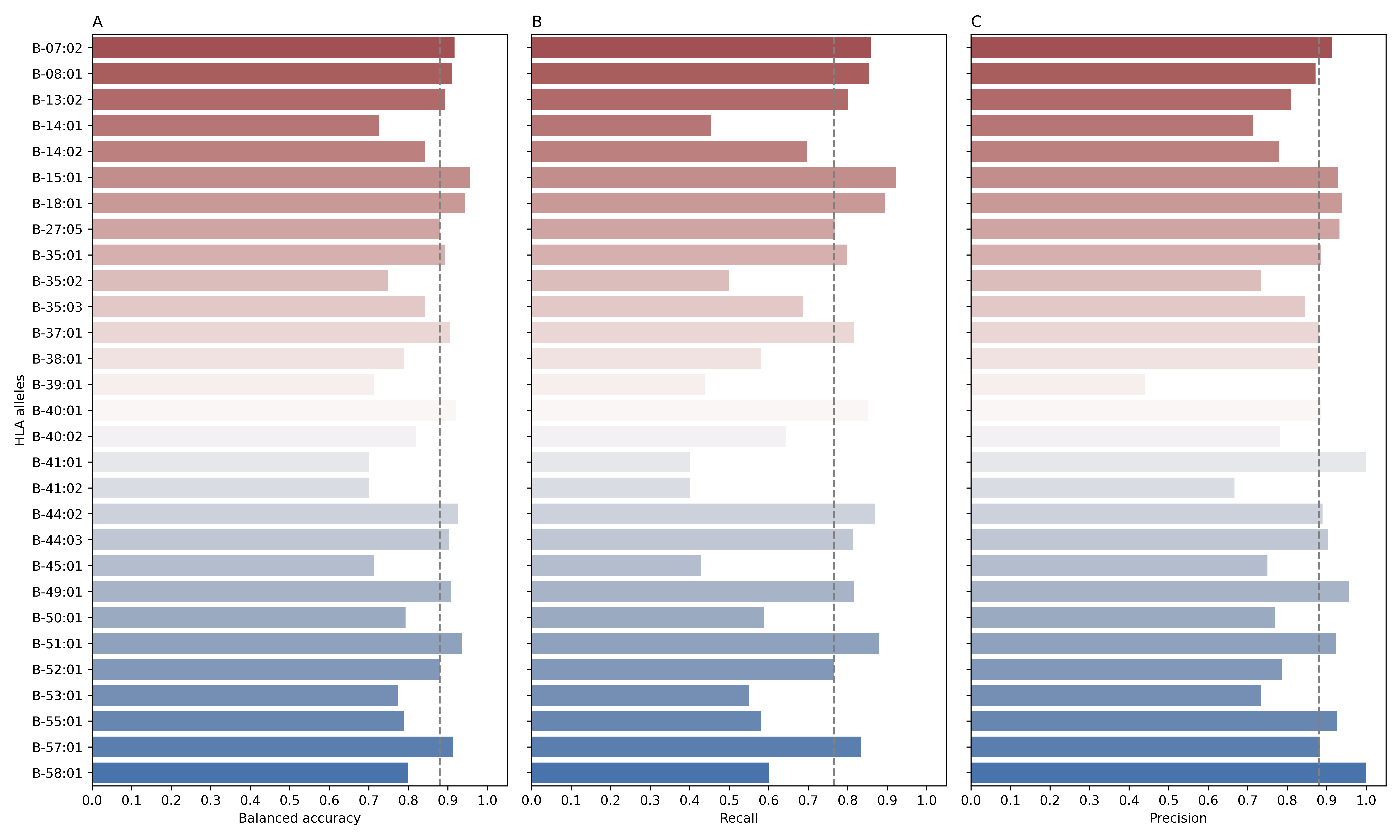

**Figure S3:** The performance of different models to predict HLA-B allele-carriership status from the TRB repertoire evaluated on a test dataset made of 1,111 repertoires. On the y-axis the models of each HLA-B allele are shown, for example, B-07:02 represents a model that predicts whether an individual is a carrier for the HLA-B*07:02 allele or not. (**A**), (**B**) and (**C**), depict the balanced accuracy, recall and precision, respectively. Across all panels, grey dashed lines represent the median.

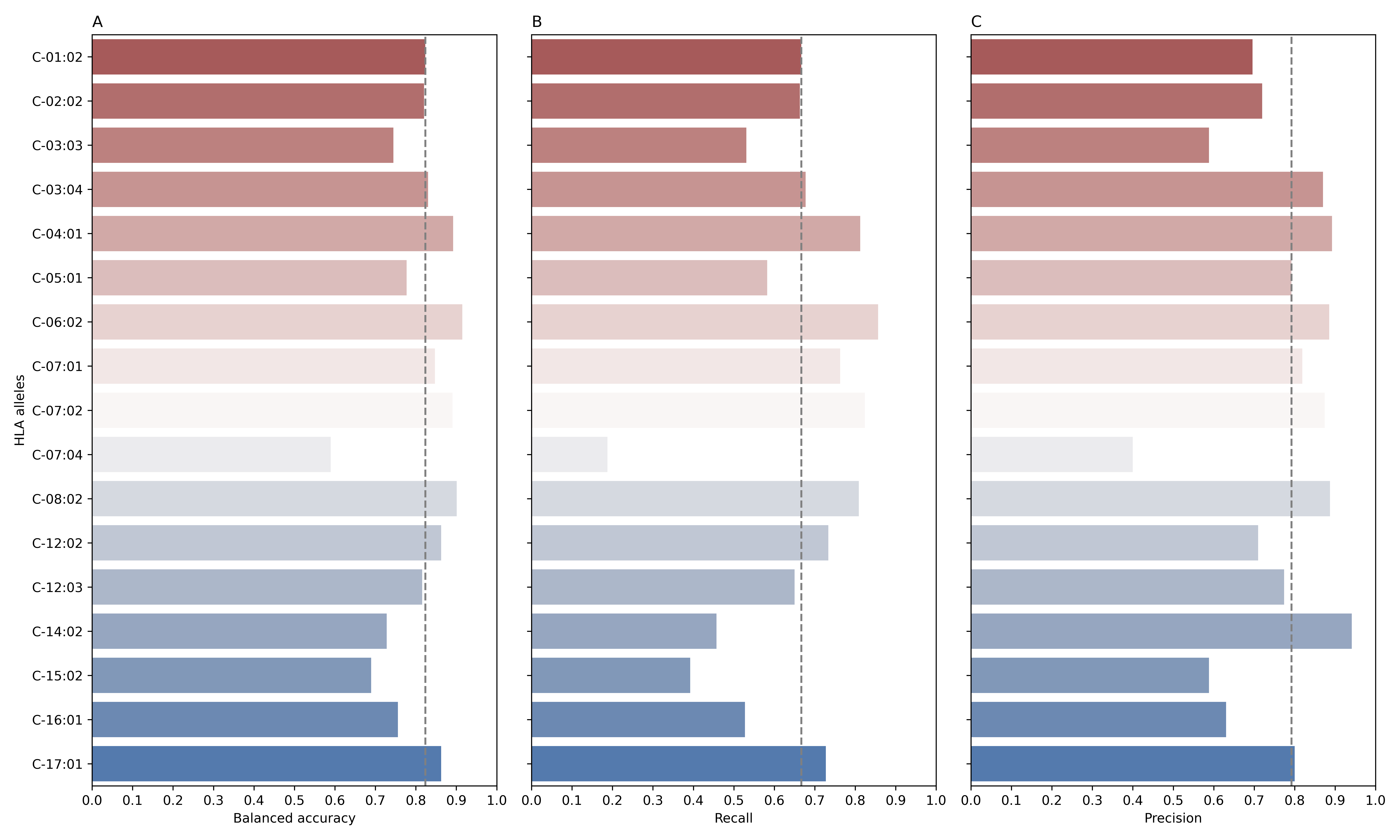

**Figure S4**: The performance of different models to predict HLA-C allele-carriership status from the TRB repertoire evaluated on a test dataset made of 1,111 repertoires. On the y-axis the models of each HLA-C allele are shown, for example, C-01:02 represents a model that predicts whether an individual is a carrier for the HLA-C*01:02 allele or not. (**A**), (**B**) and (**C**), depict the balanced accuracy, recall and precision, respectively. Across all panels, grey dashed lines represent the median.

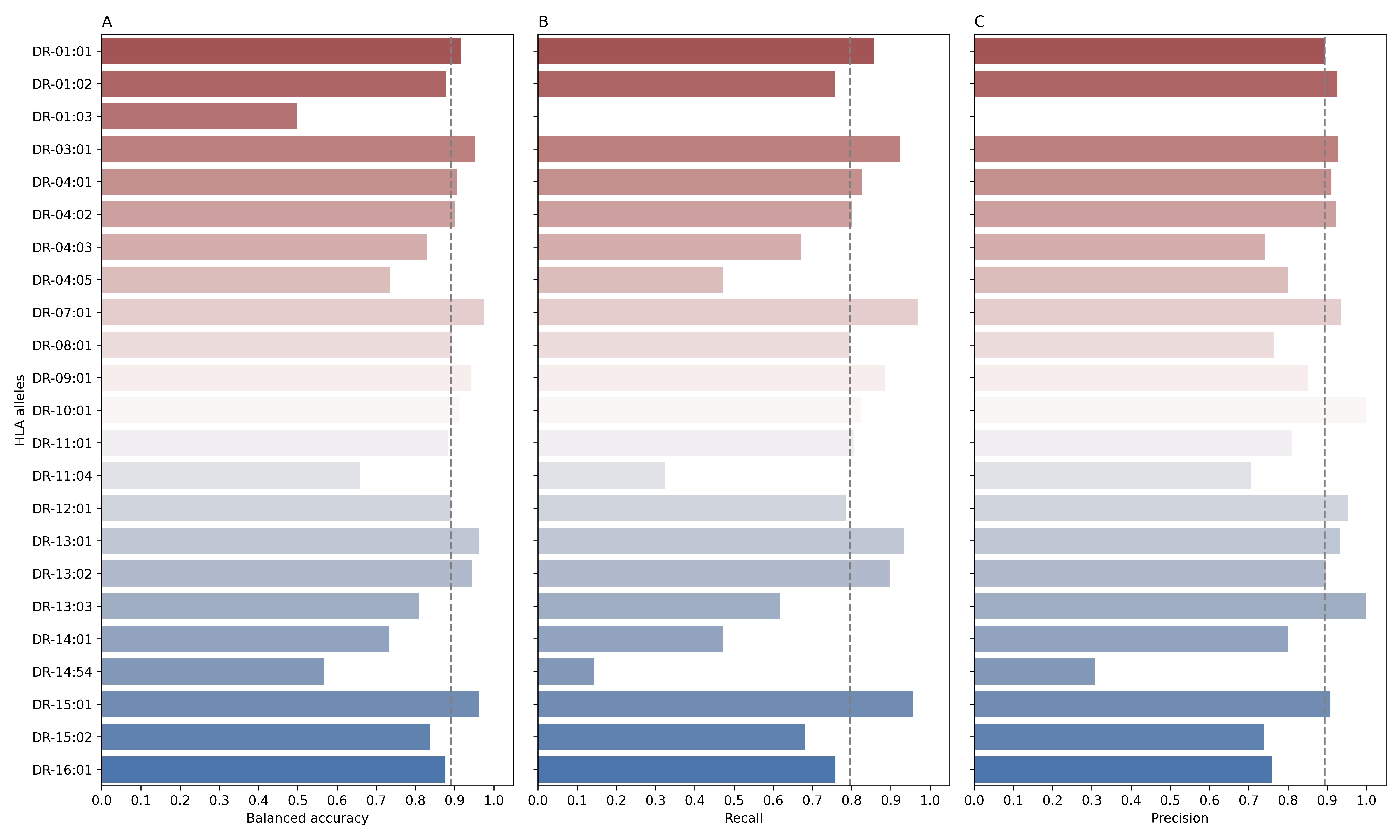

**Figure S5:** The performance of different models to predict HLA-DR allele-carriership status from the TRB repertoire evaluated on a test dataset made of 1,111 repertoires. On the y-axis the models of each HLA-DR allele are shown, for example, DR-01:01 represents a model that predicts whether an individual is a carrier for the HLA-DRB1*01:01 allele or not. (**A**), (**B**) and (**C**), depict the balanced accuracy, recall and precision, respectively. Across all panels, grey dashed lines represent the median.

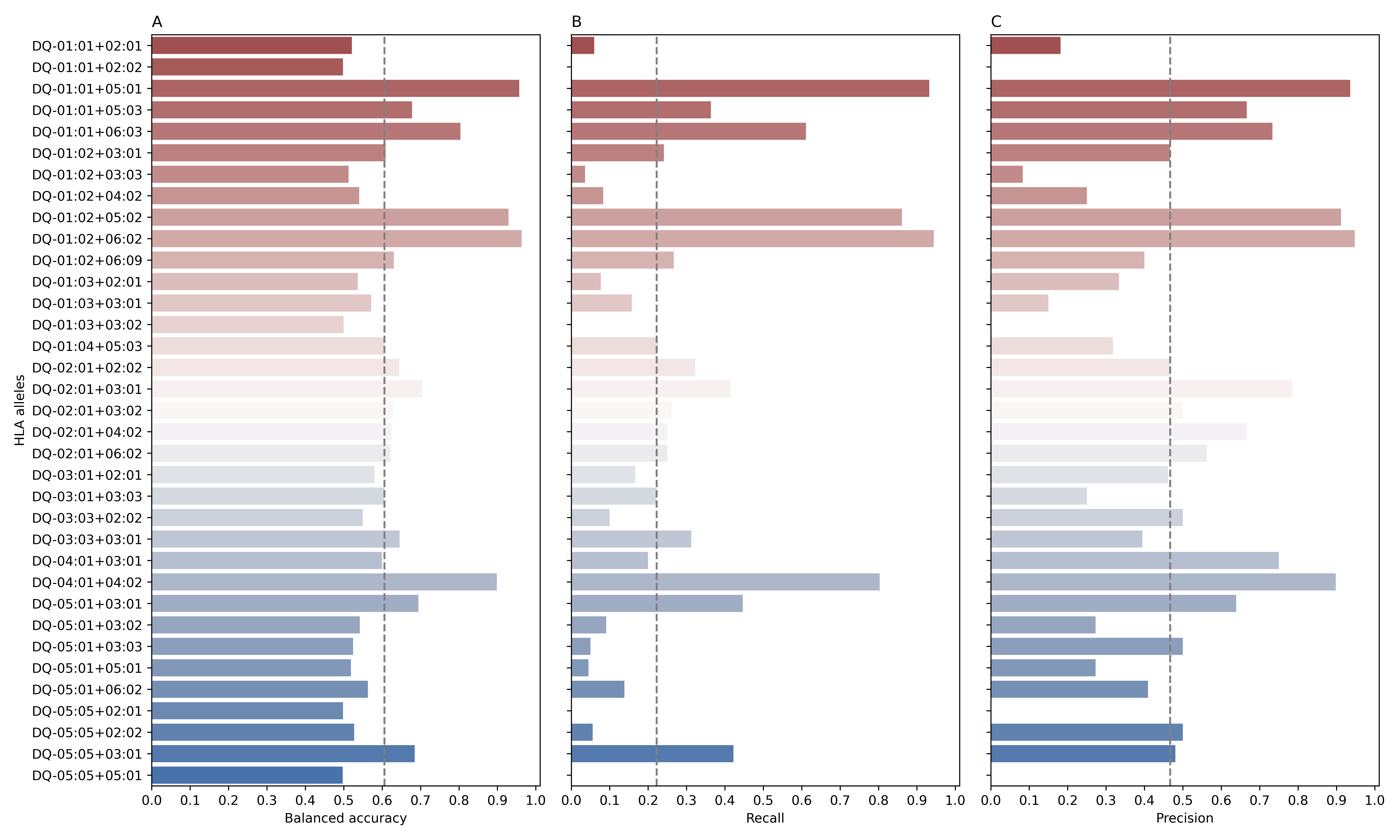

**Figure S6:** The performance of different models to predict HLA-DQ allele-carriership status from the TRB repertoire evaluated on a test dataset made of 1,111 repertoires. On the y-axis the models of each HLA-DQ allele are shown, for example, DR-01:01+02:01 represents a model that predicts whether an individual is a carrier for the HLA-DQA*01:01-DQB1*02:01 allele or not. (**A**), (**B**) and (**C**), depict the balanced accuracy, recall and precision, respectively. Across all panels, grey dashed lines represent the median.

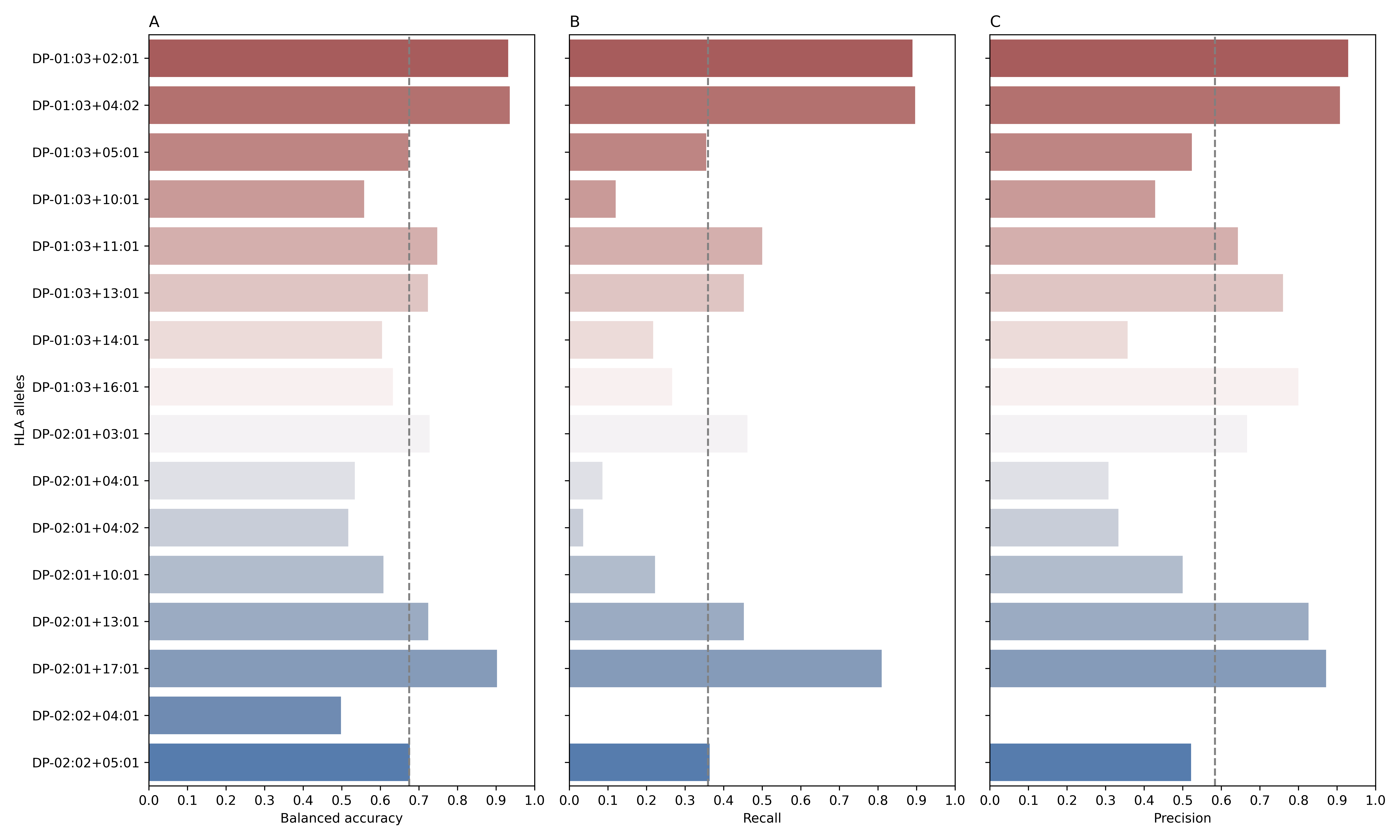

**Figure S7:** The performance of different models to predict HLA-DP allele-carriership status from the TRB repertoire evaluated on a test dataset made of 1,111 repertoires. On the y-axis the models of each HLA-DP allele are shown, for example, DP-01:03+02:01 represents a model that predicts whether an individual is a carrier for the HLA-DPA*01:03-DPB1*02:01 allele or not. (**A**), (**B**) and (**C**), depict the balanced accuracy, recall and precision, respectively. Across all panels, grey dashed lines represent the median.

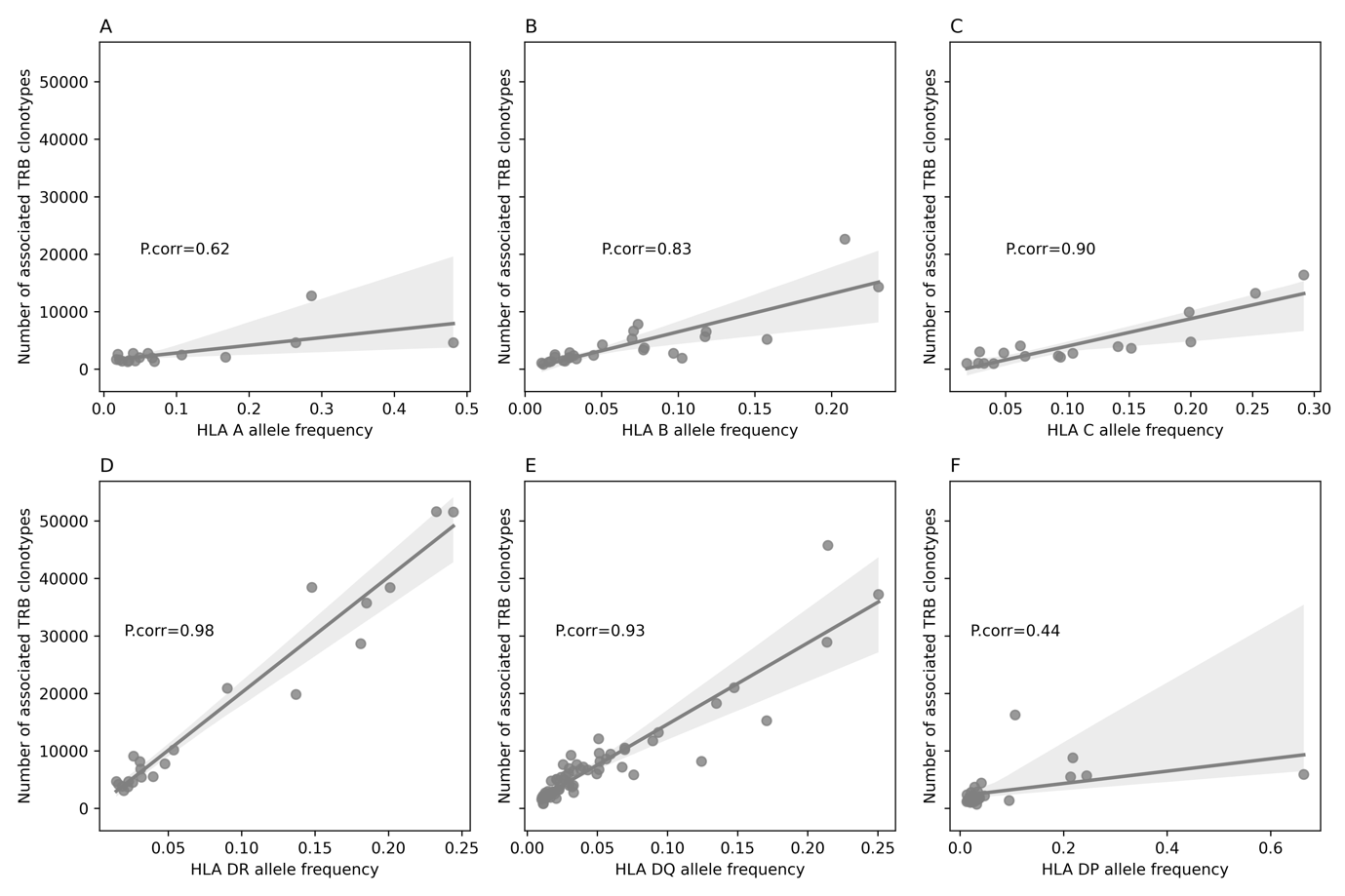

**Figure S8:** The relationship between HLA allele carriership frequency (shown on the x-axis) and the number of associated TRB clonotypes (shown on the y-axis) for the six classical HLA proteins, namely, HLA-A (**A**), HLA-B (**B**), HLA-C (**C**), HLA-DR (**D**), HLA-DQ (**E**), HLA-DP (**F**). Lastly, P.corr donates the Pearson correlation co-efficiency.

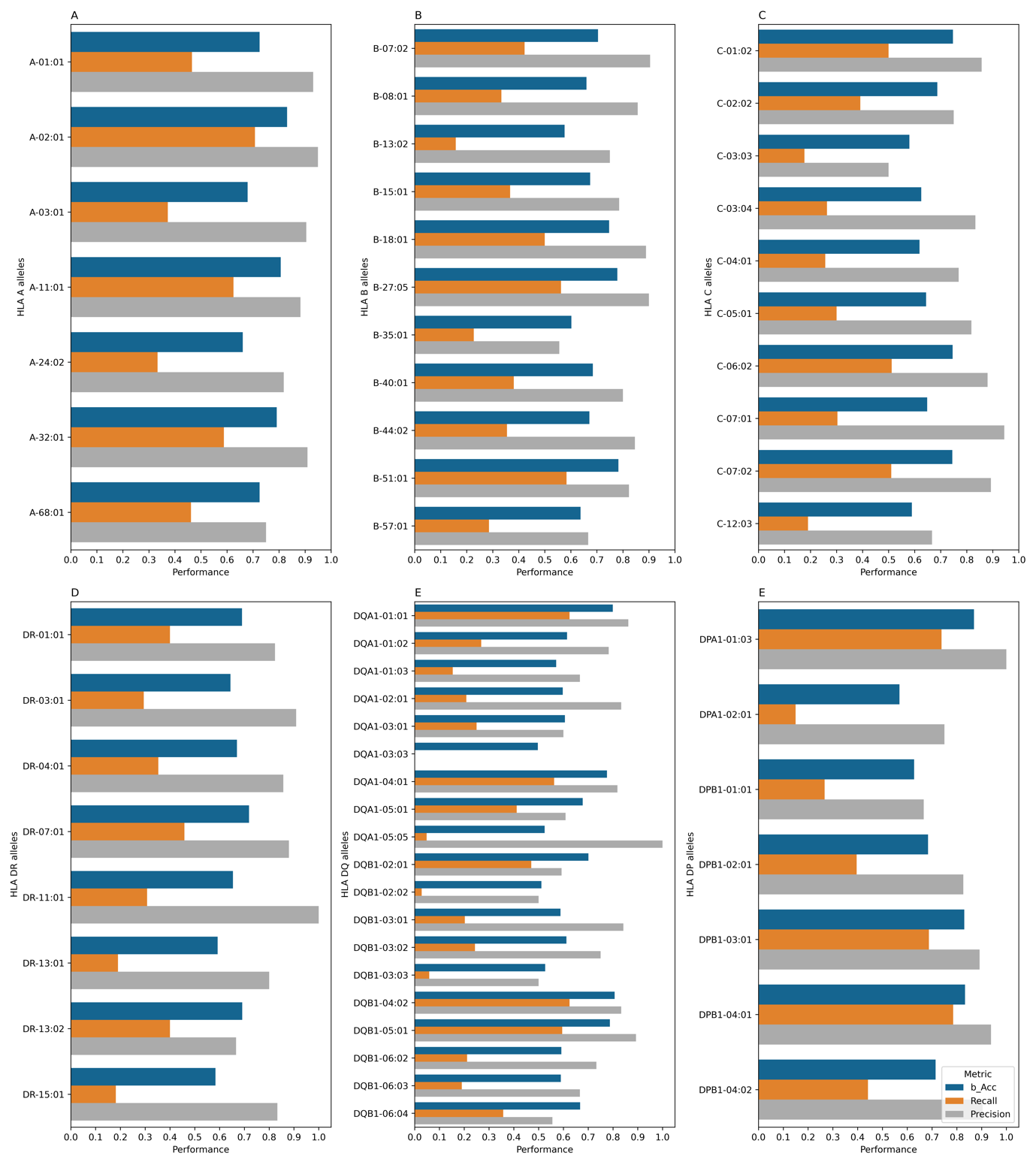

**Figure S9:** The performance of the TRB-based HLA imputation models on an independent test dataset obtained from Rosati et al.^13^. (**A**) shows the performance of HLA-A allele models, while (**B**), (**C**), and (**D**) show the performance of the HLA-B, HLA-C and HLA-DR alleles, respectively. (**E**) and (**F**) show the performance of HLA-DQA/DQB and HLA-DPA/DPB proteins, respectively. We have not performed the imputation performance at the functional αβ HLA protein level given that some of the HLA-DQ and DP genes were not typed to the same four-digit resolution. Across all panels, alleles with carriership frequency <0.05 (n<12 samples) were excluded from the analysis.

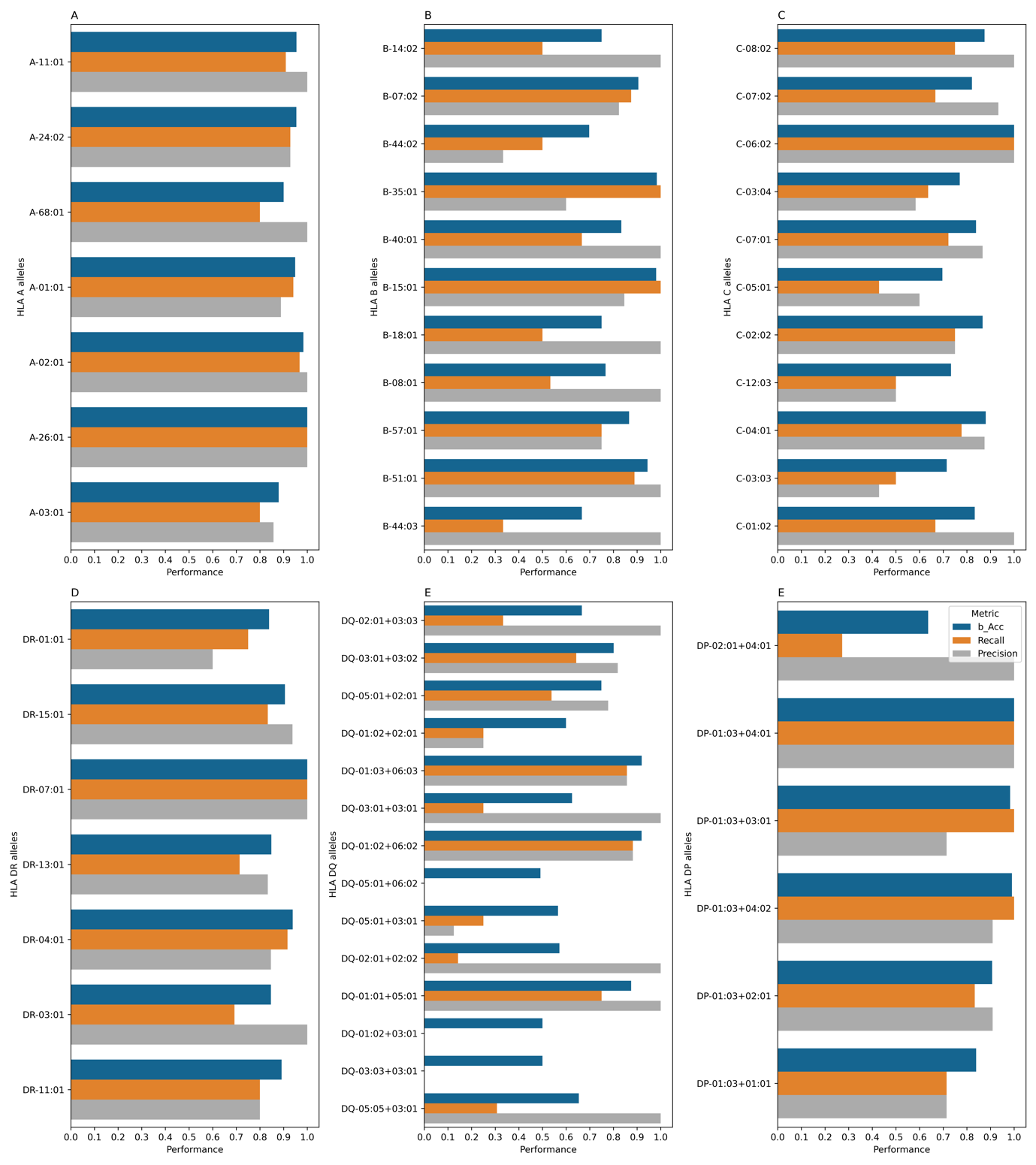

**Figure S10:** The performance of the TRB-based HLA imputation models on an independent test dataset obtained from the immuneCODE dataset^14^. (**A-E**) show the performance of HLA-A, HLA-B, HLA-C, HLA-DR, HLA-DQ and HLA-DP alleles, respectively. Across all panels, alleles with carriership frequency <0.05 (n<3 samples) were excluded from the analysis.

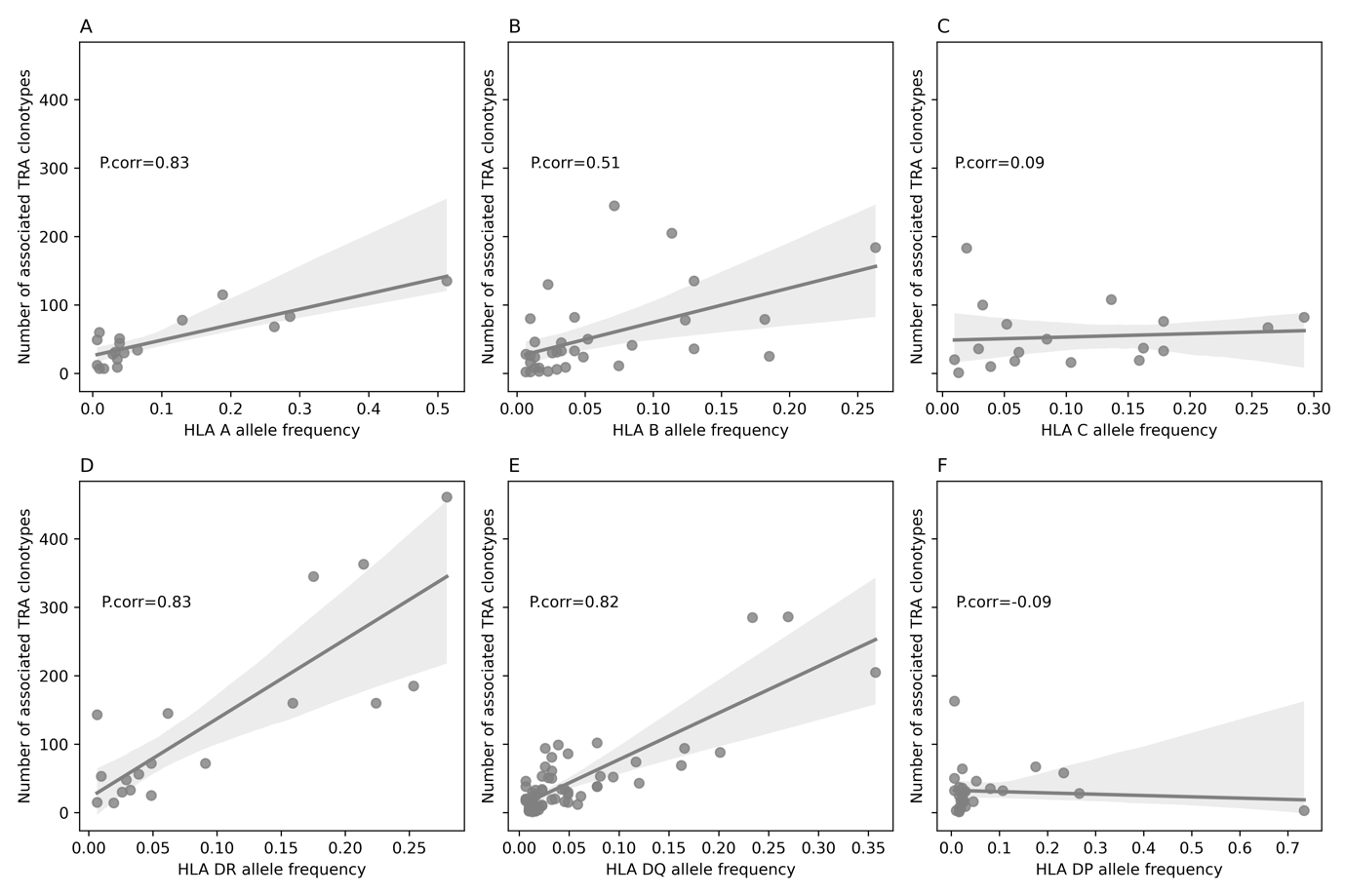

**Figure S11:** The relationship between HLA allele carriership frequency (shown on the x-axis) and the number of associated TRA clonotypes (shown on the y-axis) for the six classical HLA proteins, namely, HLA-A (**A**), HLA-B (**B**), HLA-C (**C**), HLA-DR (**D**), HLA-DQ (**E**), HLA-DP (**F**). Lastly, P.corr donates the Pearson correlation co-efficiency.

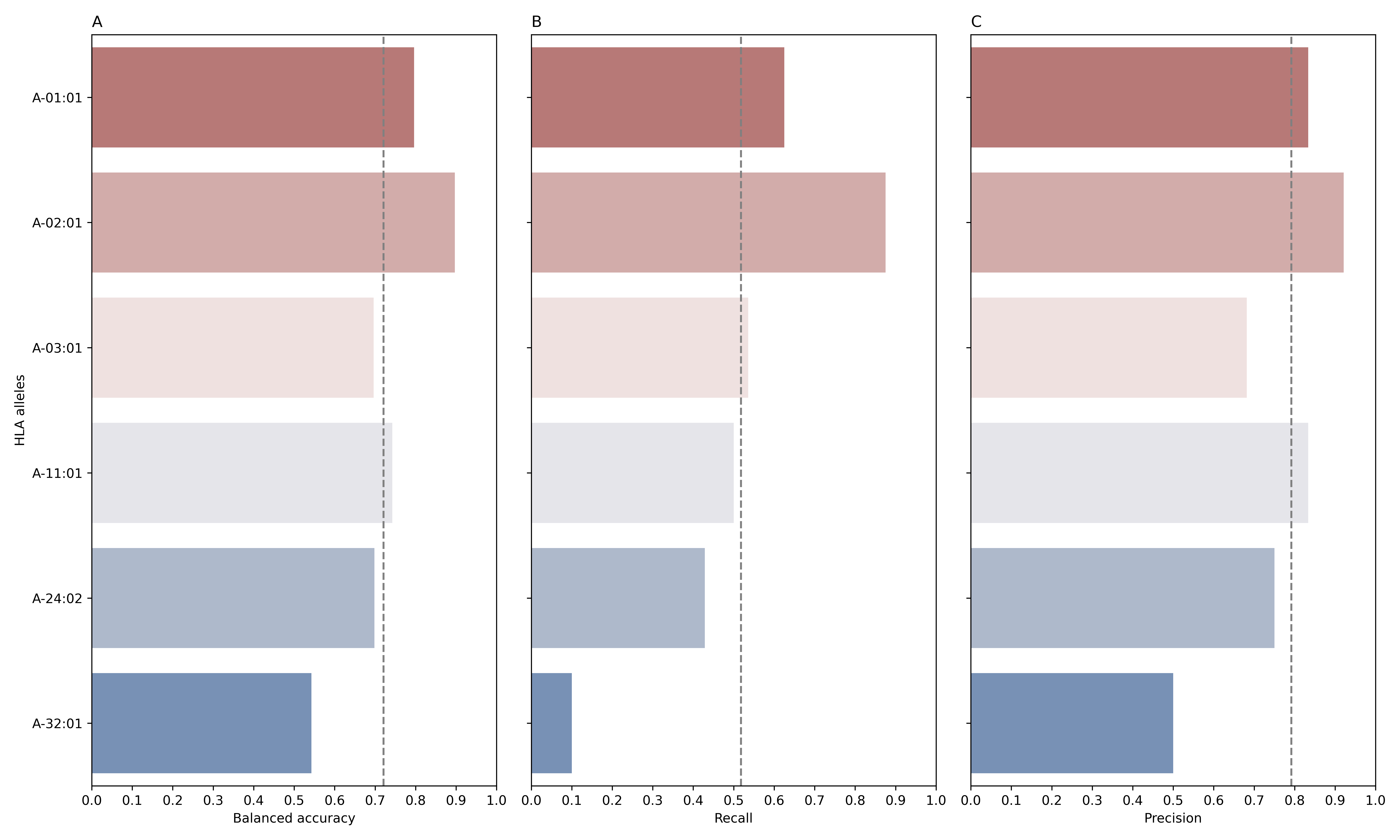

**Figure S12**: The performance of different models to predict the HLA-A allele-carriership status from the TRA repertoire evaluated on a test dataset made of 77 repertoires. These models were trained on a discovery dataset composite of 308 TRA repertoires with matching HLA calls. On the y-axis the models of each HLA-A allele are shown, for example, A-01:01 represents a model that predicts whether an individual is a carrier for the HLA-A*01:01 allele or not. (**A**), (**B**) and (**C**), depict the balanced accuracy, recall and precision, respectively. Across all panels, grey dashed lines represent the median.

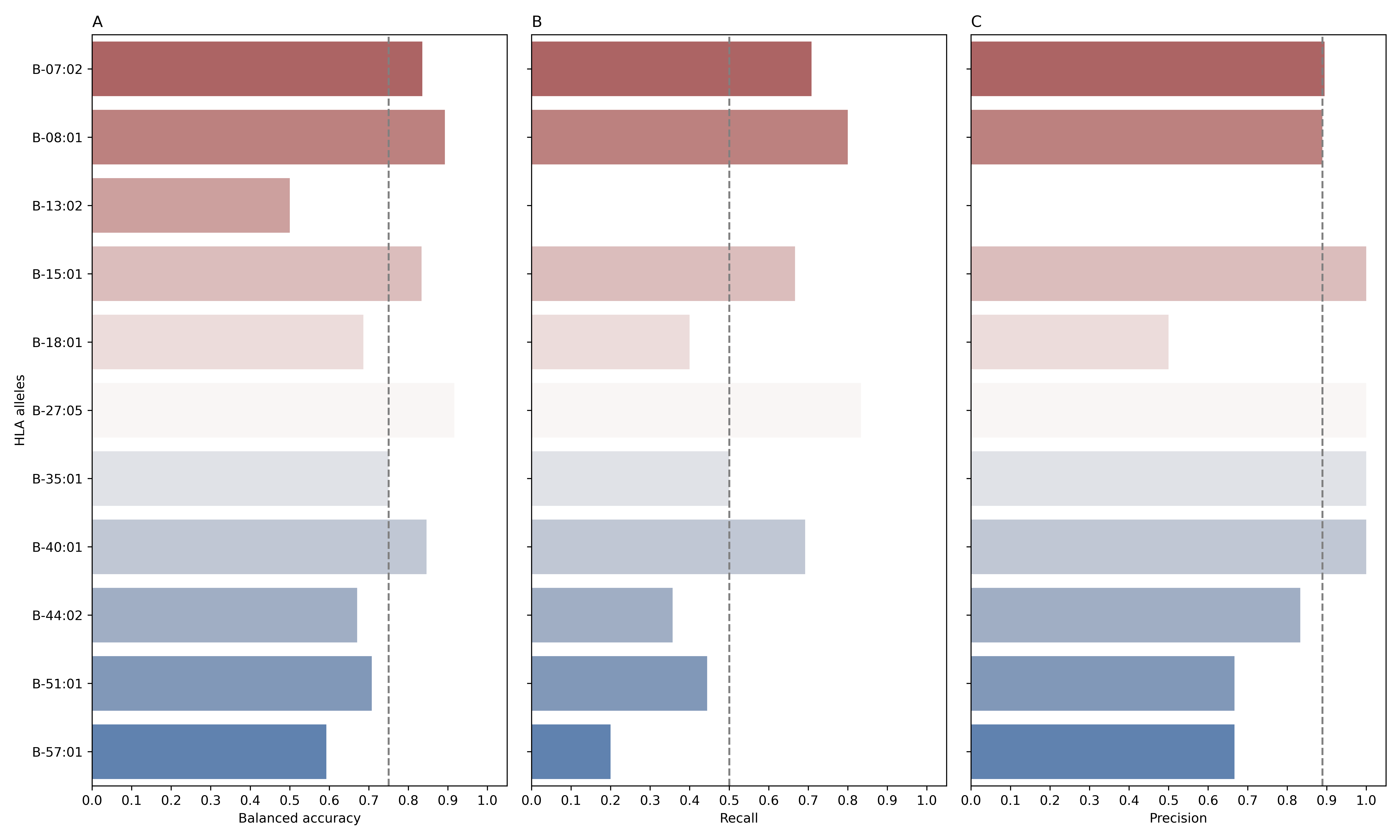

**Figure S13:** The performance of different models to predict the HLA-B allele-carriership status from the TRA repertoire evaluated on a test dataset made of 77 repertoires. These models were trained on a discovery dataset composite of 308 TRA repertoires with matching HLA calls. On the y-axis the models of each HLA-B allele are shown, for example, B*07:02 represents a model that predicts whether an individual is a carrier for the HLA-B*07:02 allele or not. (**A**), (**B**) and (**C**), depict the balanced accuracy, recall and precision, respectively. Across all panels, grey dashed lines represent the median.

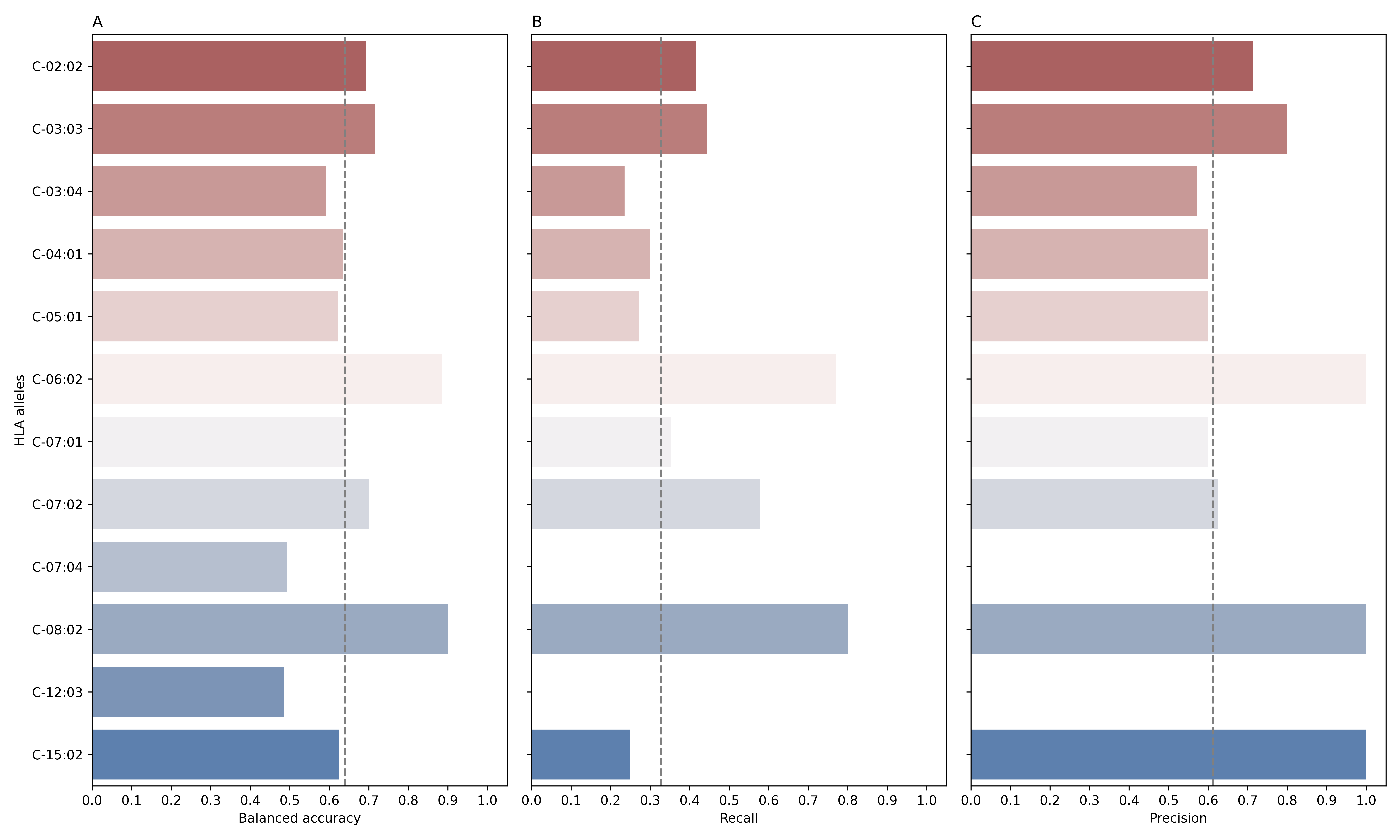

**Figure S14:** The performance of different models to predict the HLA-C allele-carriership status from the TRA repertoire evaluated on a test dataset made of 77 repertoires. These models were trained on a discovery dataset composite of 308 TRA repertoires with matching HLA calls. On the y-axis the models of each HLA-C allele are shown, for example, C*02:02 represents a model that predicts whether an individual is a carrier for the HLA-C*02:02 allele or not. (**A**), (**B**) and (**C**), depict the balanced accuracy, recall and precision, respectively. Across all panels, grey dashed lines represent the median.

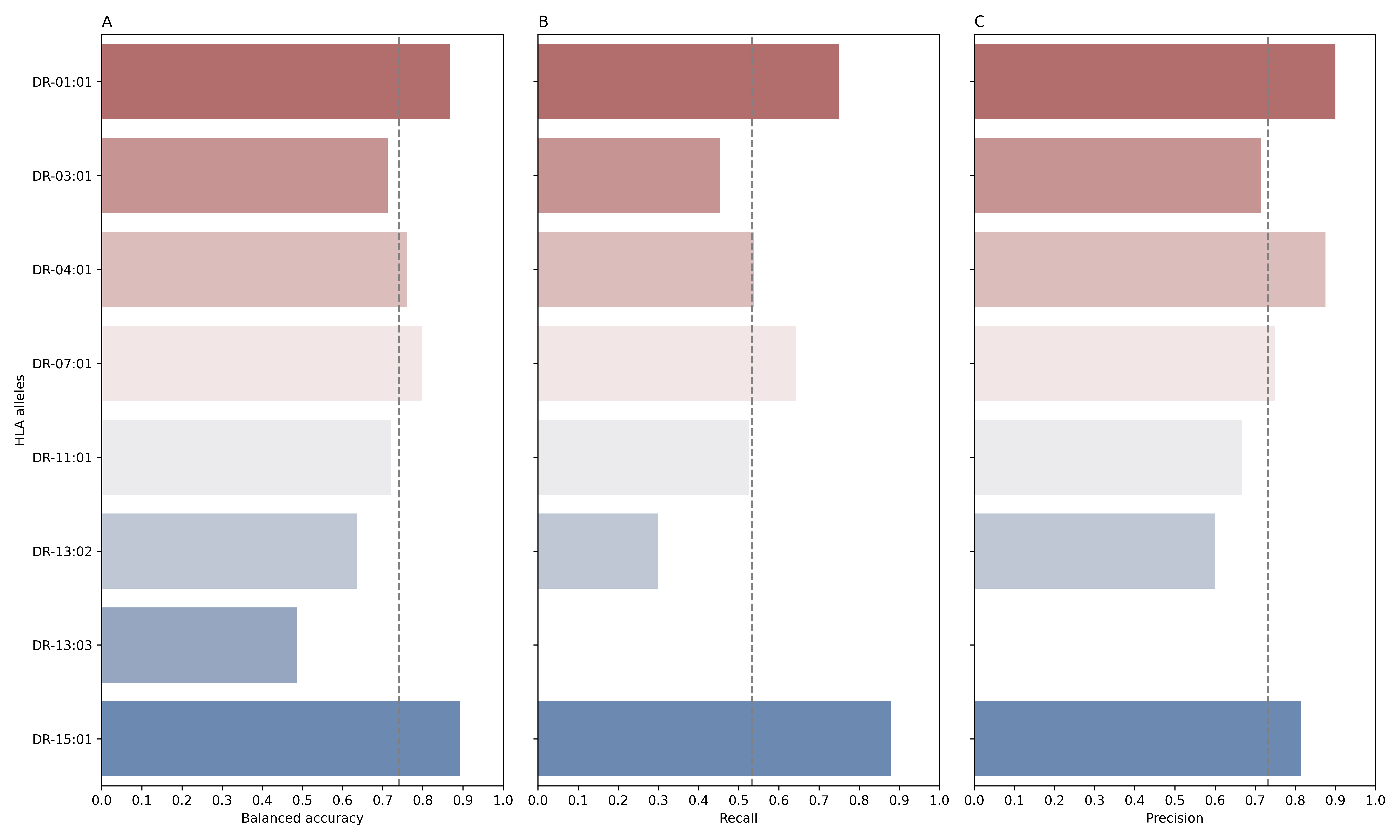

**Figure S15**: The performance of different models to predict the HLA-DR allele-carriership status from the TRA repertoire evaluated on a test dataset made of 77 repertoires. These models were trained on a discovery dataset composite of 308 TRA repertoires with matching HLA calls. On the y-axis the models of each HLA-DR allele are shown, for example, DR*01:01 represents a model that predicts whether an individual is a carrier for the HLA-DRB1*01:01 allele or not. (**A**), (**B**) and (**C**), depict the balanced accuracy, recall and precision, respectively. Across all panels, grey dashed lines represent the median.

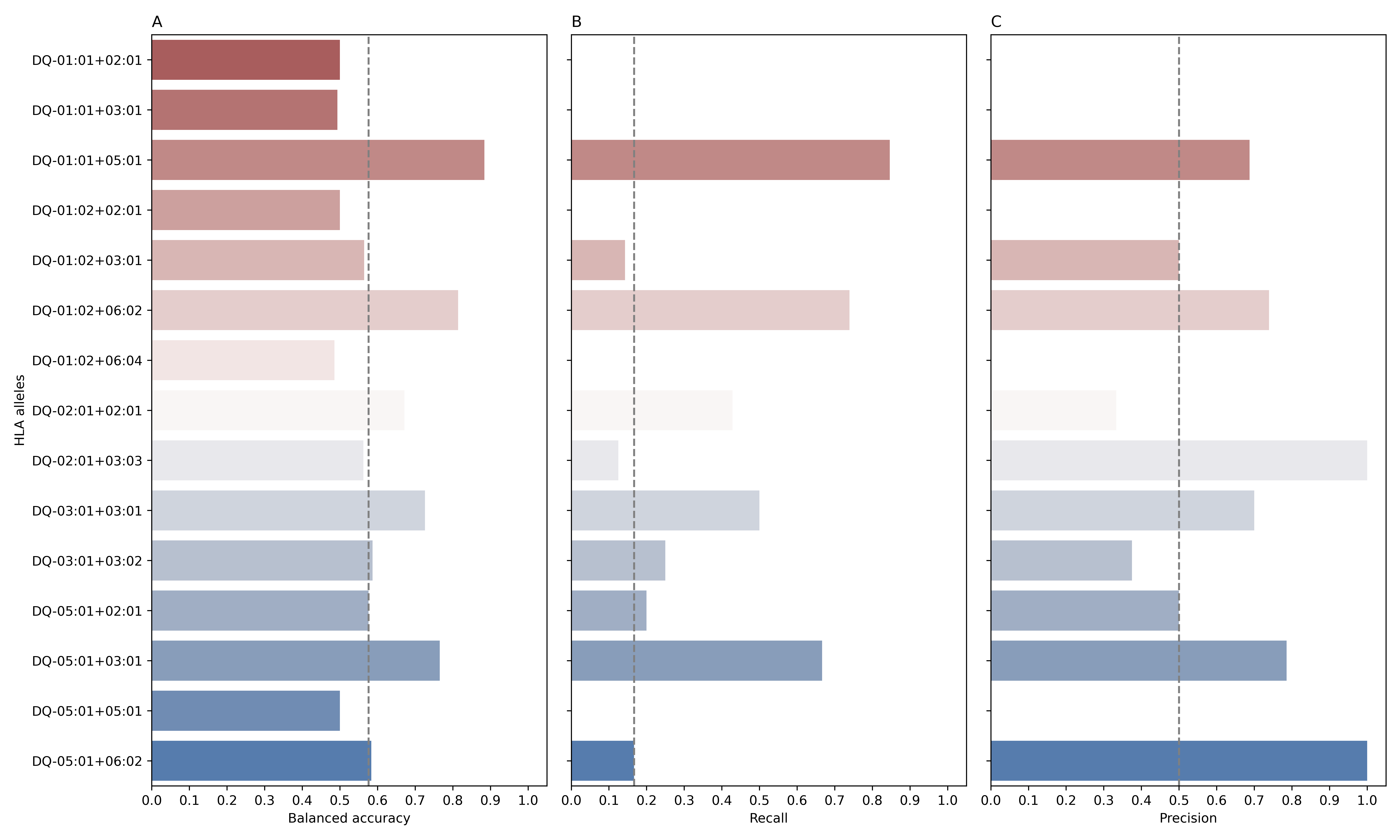

**Figure S16:** The performance of different models to predict the HLA-DQ allele-carriership status from the TRA repertoire evaluated on a test dataset made of 77 repertoires. These models were trained on a discovery dataset composite of 308 TRA repertoires with matching HLA calls. On the y-axis the models of each HLA-DQ allele are shown, for example, DQ*01:01+02:01 represents a model that predicts whether an individual is a carrier for the HLA-DQA*01:01-DQB1*02:01 allele or not. (**A**), (**B**) and (**C**), depict the balanced accuracy, recall and precision, respectively. Across all panels, grey dashed lines represent the median.

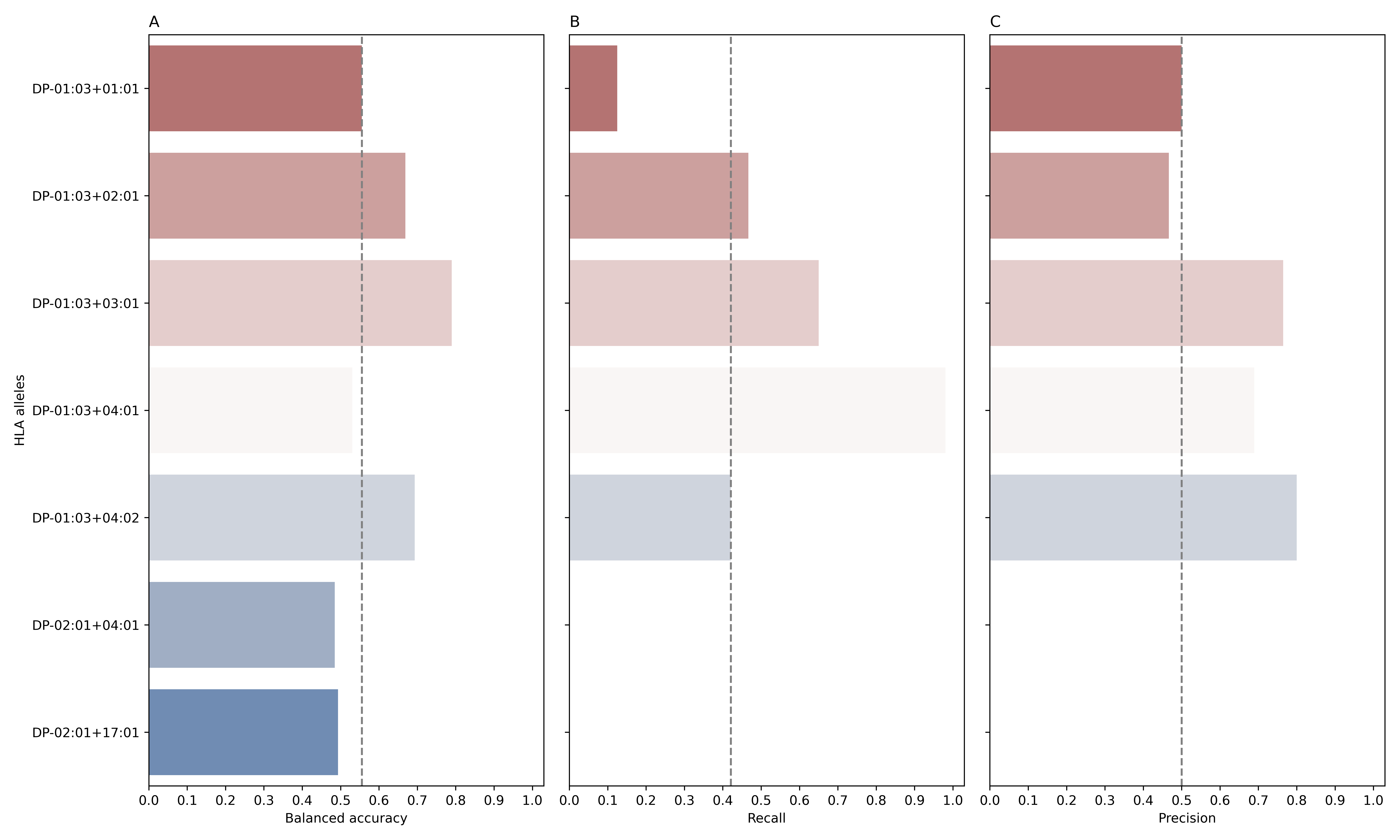

**Figure S17:** The performance of different models to predict the HLA-DP allele-carriership status from the TRA repertoire evaluated on a test dataset made of 77 repertoires. These models were trained on a discovery dataset composite of 308 TRA repertoires with matching HLA calls. On the y-axis the models of each HLA-DP allele are shown, for example, DPA*01:03+01:01 represents a model that predicts whether an individual is a carrier for the HLA-DPA*03:01-DPB1*01:01 allele or not. (**A**), (**B**) and (**C**), depict the balanced accuracy, recall and precision, respectively. Across all panels, grey dashed lines represent the median.

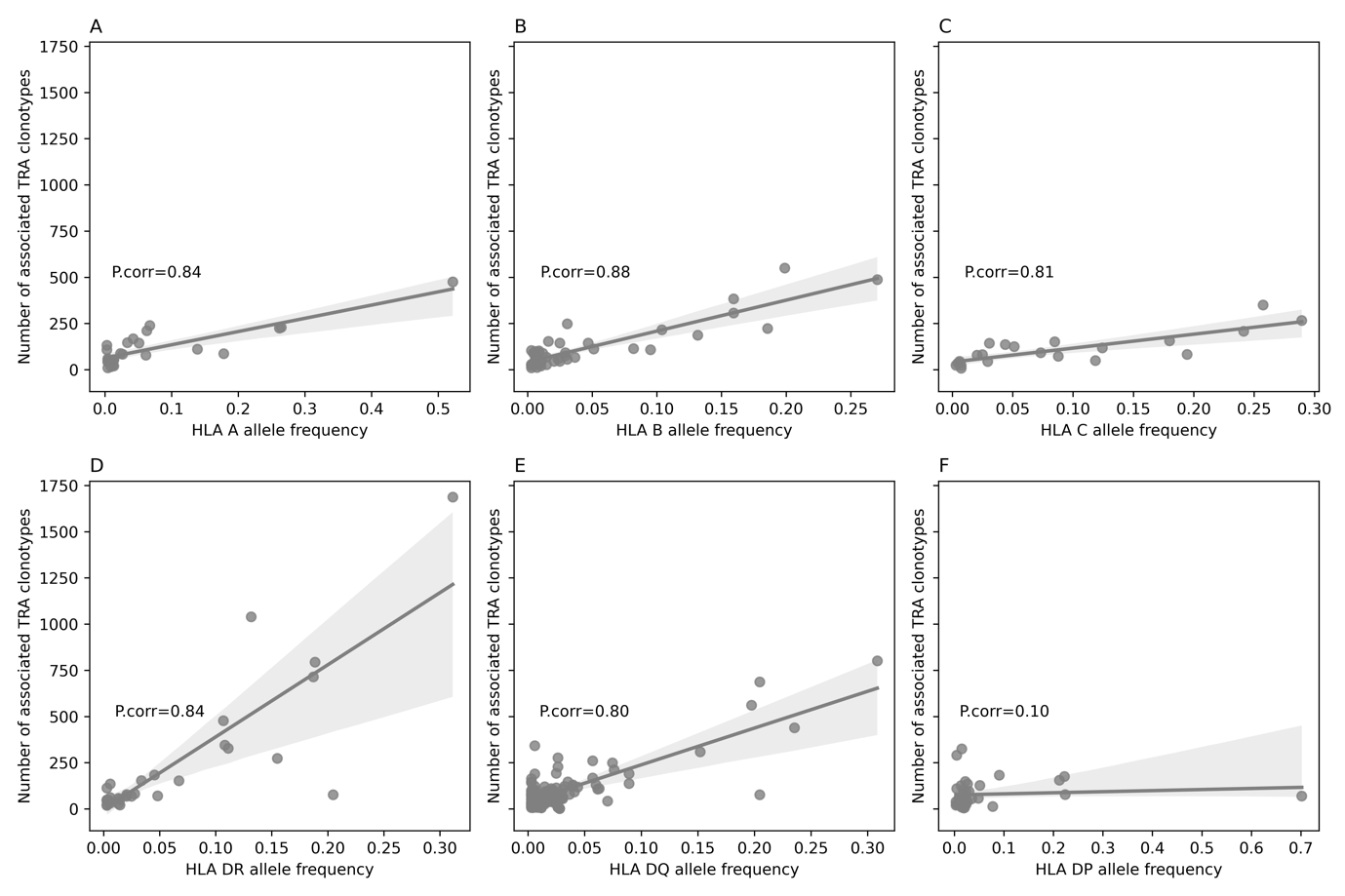

**Figure S18:** The relationship between HLA allele carriership frequency (shown on the x-axis) and the number of associated TRA clonotypes (shown on the y-axis) for the six classical HLA proteins, namely, HLA-A (**A**), HLA-B (**B**), HLA-C (**C**), HLA-DR (**D**), HLA-DQ (**E**), HLA-DP (**F**). Lastly, P.corr donates the Pearson correlation co-efficiency.

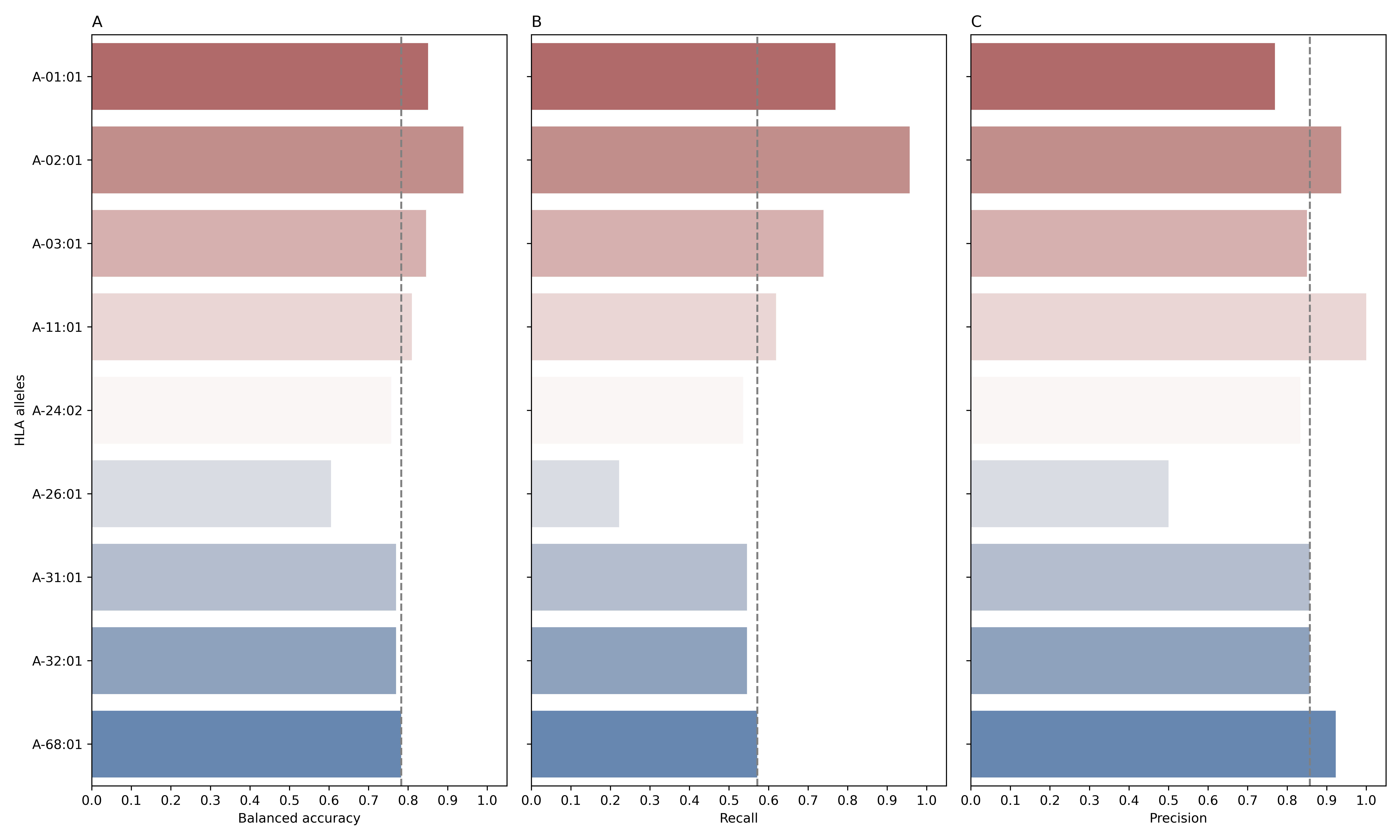

**Figure S19**: The performance of different models to predict the HLA-A allele-carriership status from the TRA repertoire evaluated on a test dataset made of 171repertoires. These models were trained on a discovery dataset composite of 684 TRA repertoires with matching HLA calls. On the y-axis the models of each HLA-A allele are shown, for example, A-01:01 represents a model that predicts whether an individual is a carrier for the HLA-A*01:01 allele or not. (**A**), (**B**) and (**C**), depict the balanced accuracy, recall and precision, respectively. Across all panels, grey dashed lines represent the median.

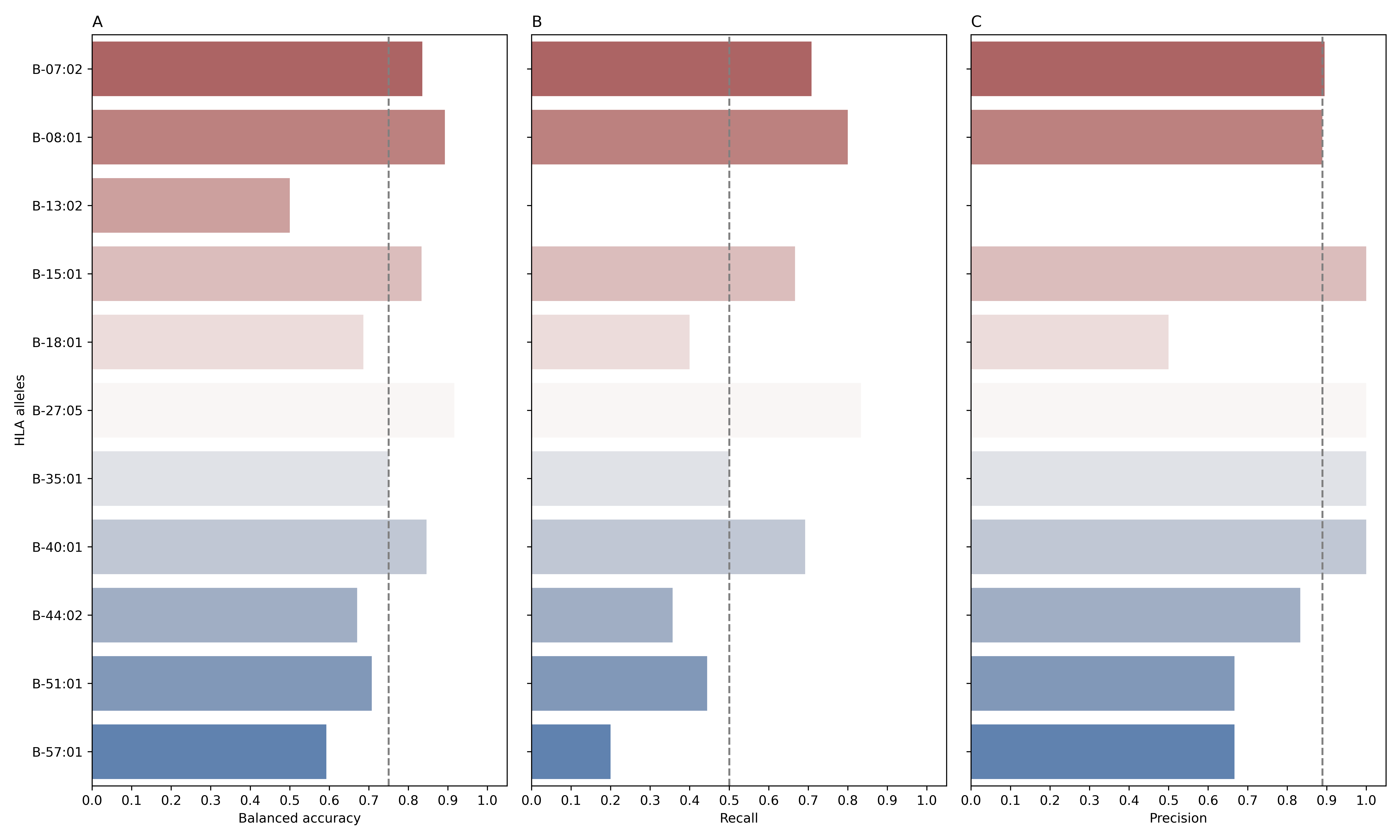

**Figure S20:** The performance of different models to predict the HLA-B allele-carriership status from the TRA repertoire evaluated on a test dataset made of 171 repertoires. These models were trained on a discovery dataset composite of 684 TRA repertoires with matching HLA calls. On the y-axis the models of each HLA-B allele are shown, for example, B-07:02 represents a model that predicts whether an individual is a carrier for the HLA-B*07:02 allele or not. (**A**), (**B**) and (**C**), depict the balanced accuracy, recall and precision, respectively. Across all panels, grey dashed lines represent the median.

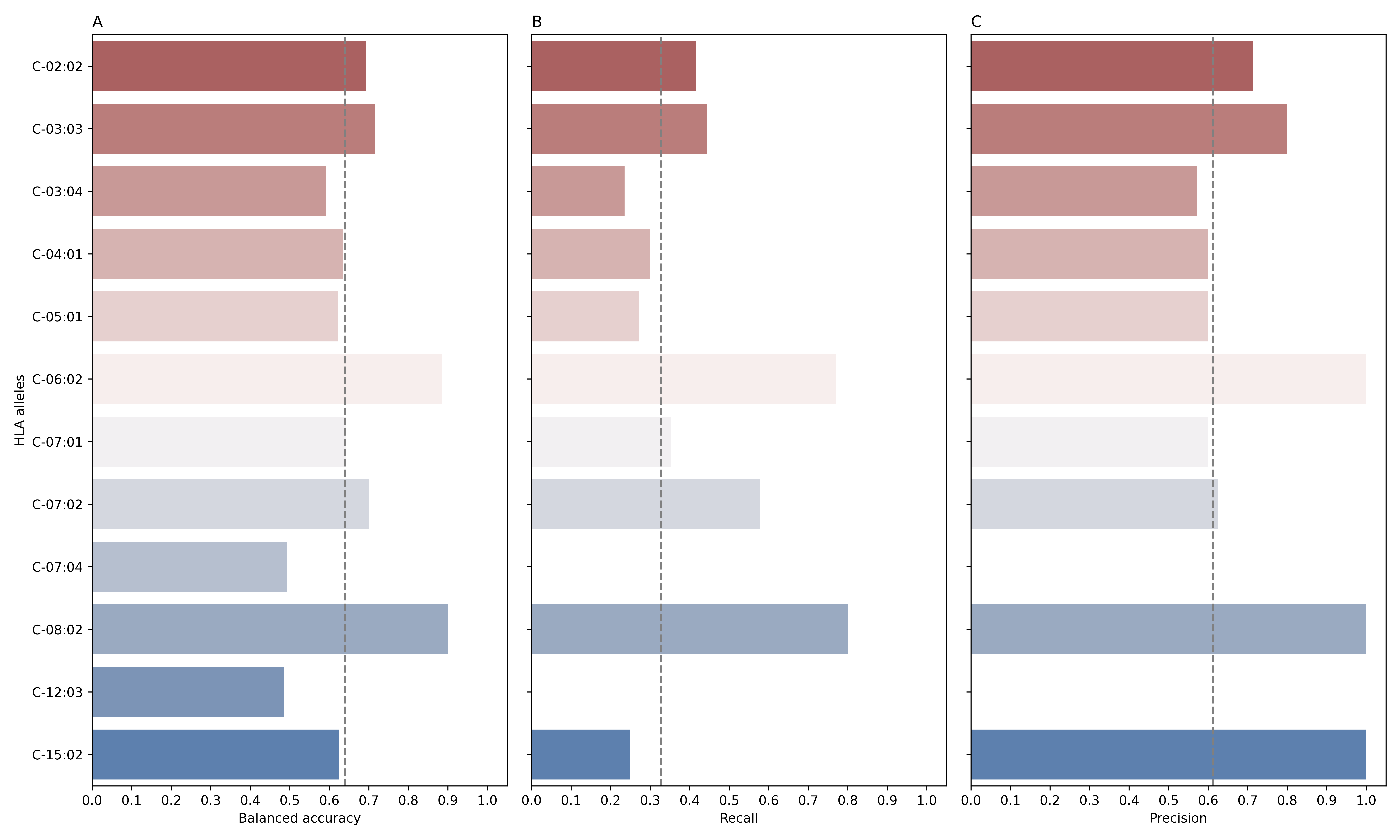

**Figure S21:** The performance of different models to predict the HLA-C allele-carriership status from the TRA repertoire evaluated on a test dataset made of 171 repertoires. These models were trained on a discovery dataset composite of 684 TRA repertoires with matching HLA calls. On the y-axis the models of each HLA-C allele are shown, for example, C-02:02 represents a model that predicts whether an individual is a carrier for the HLA-C*02:02 allele or not. (**A**), (**B**) and (**C**), depict the balanced accuracy, recall and precision, respectively. Across all panels, grey dashed lines represent the median.

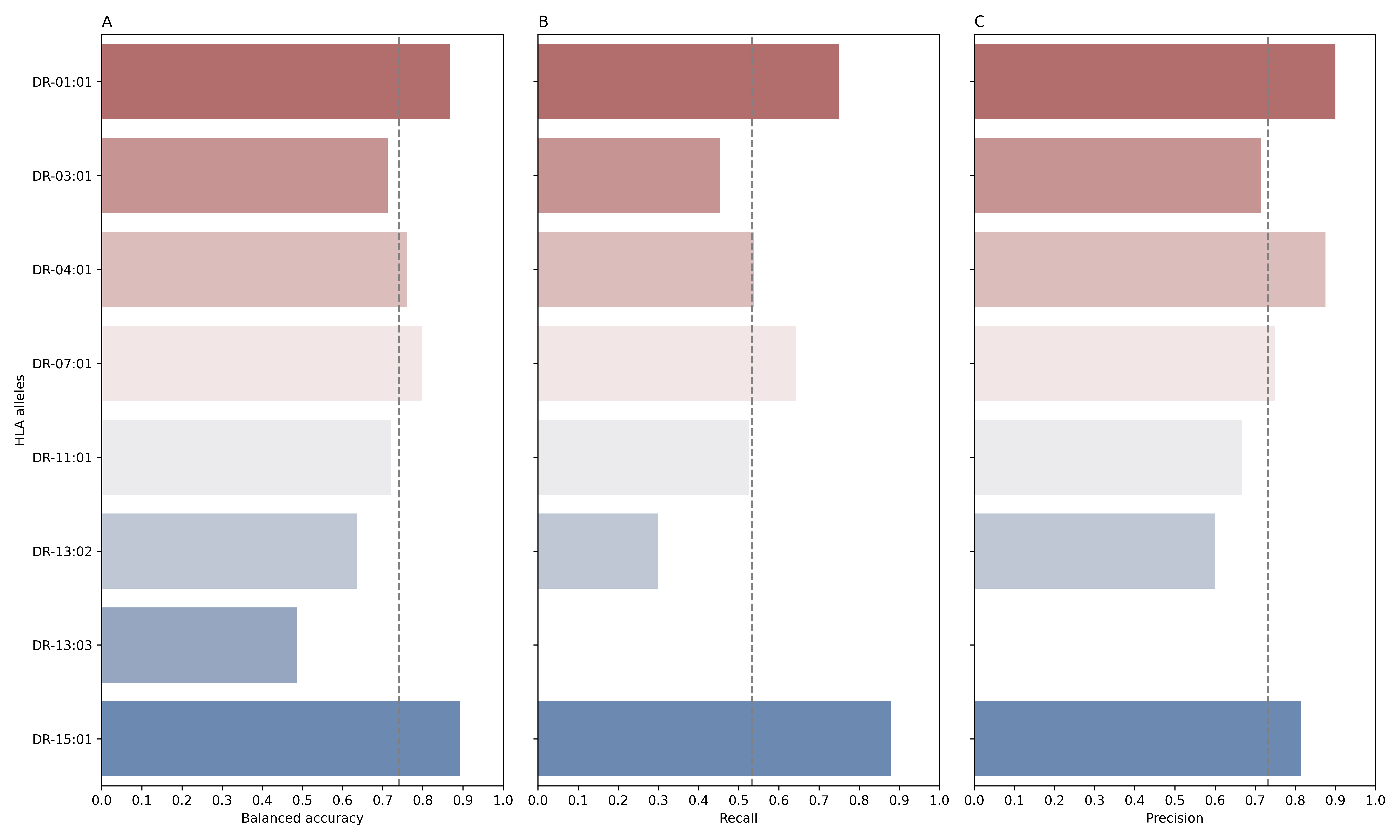

**Figure S22:** The performance of different models to predict the HLA-DR allele-carriership status from the TRA repertoire evaluated on a test dataset made of 171 repertoires. These models were trained on a discovery dataset composite of 684 TRA repertoires with matching HLA calls. On the y-axis the models of each HLA-DR allele are shown, for example, DR-01:01 represents a model that predicts whether an individual is a carrier for the HLA-DRB1*01:01 allele or not. (**A**), (**B**) and (**C**), depict the balanced accuracy, recall and precision, respectively. Across all panels, grey dashed lines represent the median.

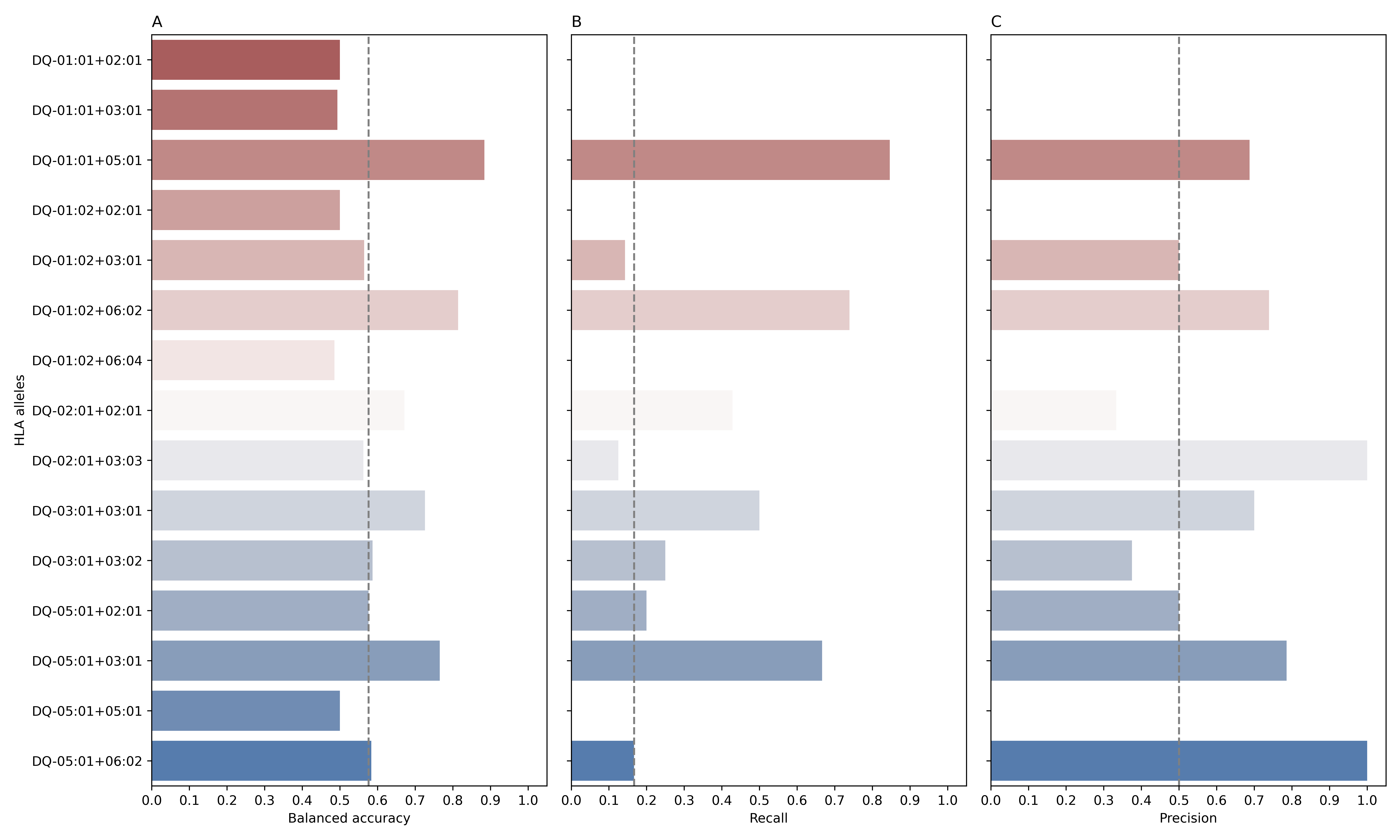

**Figure S23:** The performance of different models to predict the HLA-DQ allele-carriership status from the TRA repertoire evaluated on a test dataset made of 171 repertoires. These models were trained on a discovery dataset composite of 684 TRA repertoires with matching HLA calls. On the y-axis the models of each HLA-DQ allele are shown, for example, DQ-01:01+02:01 represents a model that predicts whether an individual is a carrier for the HLA-DQA1*01:01-DQB1*02:01 allele or not. (**A**), (**B**) and (**C**), depict the balanced accuracy, recall and precision, respectively. Across all panels, grey dashed lines represent the median.

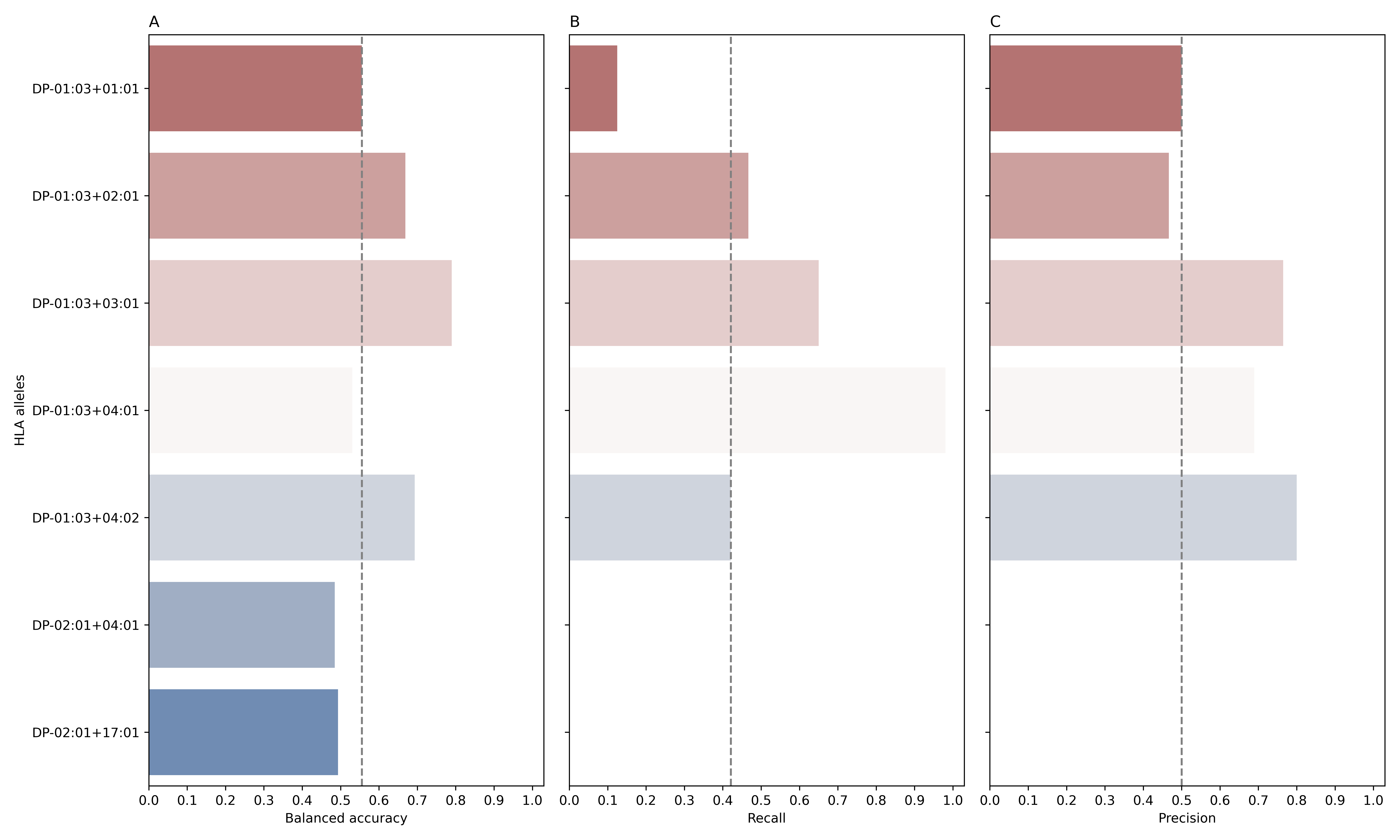

**Figure S24:** The performance of different models to predict the HLA-DP allele-carriership status from the TRA repertoire evaluated on a test dataset made of 171 repertoires. These models were trained on a discovery dataset composite of 684 TRA repertoires with matching HLA calls. On the y-axis the models of each HLA-DP allele are shown, for example, DP-01:03+01:01 represents a model that predicts whether an individual is a carrier for the HLA-DPA1*01:03-DPB1*01:01 allele or not. (**A**), (**B**) and (**C**), depict the balanced accuracy, recall and precision, respectively. Across all panels, grey dashed lines represent the median.

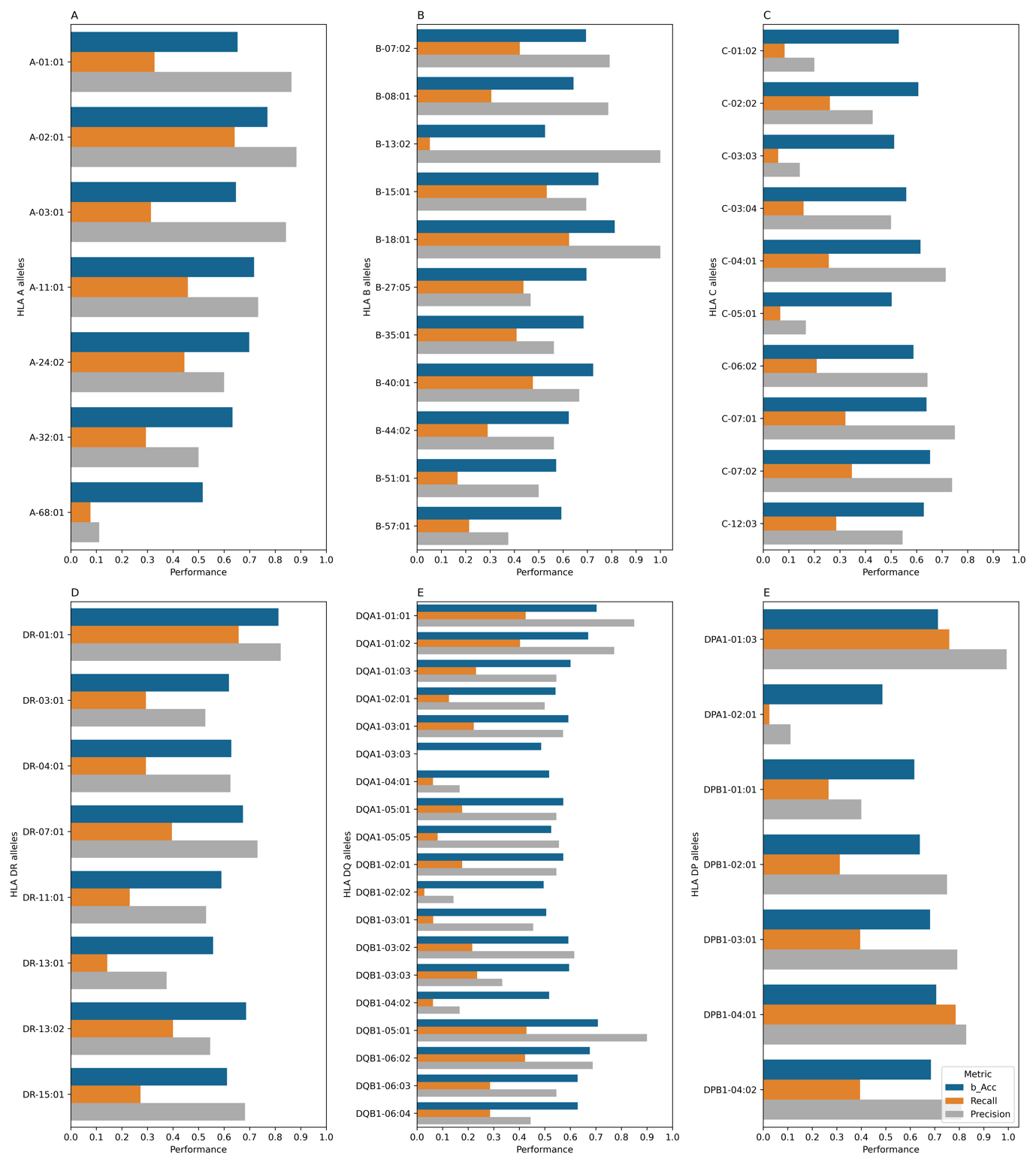

**Figure 26:** The performance of the developed TRA-based imputation models on a test dataset of paired TRA repertoire and HLA alleles that was generated by Rosati et al.^13^. (**A-E**) shows the performance on the HLA-A, HLA-B, HLA-C, HLA-DR, HLA-DQA/DQB and HLA-DPA/DPB alleles, respectively. Across all panels, alleles with carriership frequency <0.05 (n<12 samples) were excluded from the analysis.

**Supplementary tables**

**Table S1**: The number of associated TRB clonotypes per each HLA allele. The first column donates the name of the HLA protein/loci, the allele name at the two fields or four digits resolution which is equivalent to the P-group definition. The last column represents the number of associated TRB per each HLA allele.

| HLA protein | HLA allele | Number of associated clonotypes |
| --- | --- | --- |
| A | 01:01 | 12765 |
| A | 02:01 | 4612 |
| A | 02:05 | 1626 |
| A | 03:01 | 4602 |
| A | 11:01 | 2463 |
| A | 23:01 | 1516 |
| A | 24:02 | 2083 |
| A | 25:01 | 1303 |
| A | 26:01 | 1961 |
| A | 29:02 | 2026 |
| A | 30:01 | 2769 |
| A | 30:02 | 1704 |
| A | 31:01 | 1431 |
| A | 32:01 | 1330 |
| A | 33:01 | 2608 |
| A | 68:01 | 2771 |
| A | 68:02 | 1401 |
| B | 07:02 | 14331 |
| B | 08:01 | 22624 |
| B | 13:02 | 6662 |
| B | 14:01 | 841 |
| B | 14:02 | 4270 |
| B | 15:01 | 6532 |
| B | 18:01 | 3359 |
| B | 27:05 | 3706 |
| B | 35:01 | 5642 |
| B | 35:02 | 2545 |
| B | 35:03 | 2033 |
| B | 37:01 | 1498 |
| B | 38:01 | 2425 |
| B | 39:01 | 1404 |
| B | 40:01 | 2759 |
| B | 40:02 | 2000 |
| B | 44:02 | 5203 |
| B | 44:03 | 7838 |
| B | 45:01 | 1094 |
| B | 49:01 | 2889 |
| B | 50:01 | 2105 |
| B | 51:01 | 1930 |
| B | 52:01 | 2372 |
| B | 53:01 | 1254 |
| B | 55:01 | 1804 |
| B | 57:01 | 5307 |
| B | 58:01 | 1360 |
| C | 01:02 | 2247 |
| C | 02:02 | 2087 |
| C | 03:03 | 2752 |
| C | 03:04 | 3947 |
| C | 04:01 | 4736 |
| C | 05:01 | 3633 |
| C | 06:02 | 9925 |
| C | 07:01 | 16409 |
| C | 07:02 | 13220 |
| C | 07:04 | 1021 |
| C | 08:02 | 4051 |
| C | 12:02 | 3029 |
| C | 12:03 | 2316 |
| C | 14:02 | 1054 |
| C | 15:02 | 976 |
| C | 16:01 | 2820 |
| C | 17:01 | 1014 |
| DP | 01:03+01:01 | 16270 |
| DP | 01:03+02:01 | 5674 |
| DP | 01:03+03:01 | 8789 |
| DP | 01:03+04:01 | 5926 |
| DP | 01:03+04:02 | 5506 |
| DP | 01:03+05:01 | 1797 |
| DP | 01:03+09:01 | 1207 |
| DP | 01:03+10:01 | 1962 |
| DP | 01:03+11:01 | 2785 |
| DP | 01:03+13:01 | 1549 |
| DP | 01:03+14:01 | 1043 |
| DP | 01:03+15:01 | 1388 |
| DP | 01:03+16:01 | 1188 |
| DP | 01:03+17:01 | 2905 |
| DP | 01:03+19:01 | 2367 |
| DP | 02:01+01:01 | 2192 |
| DP | 02:01+02:01 | 739 |
| DP | 02:01+03:01 | 1727 |
| DP | 02:01+04:01 | 1385 |
| DP | 02:01+04:02 | 1183 |
| DP | 02:01+05:01 | 1699 |
| DP | 02:01+09:01 | 1331 |
| DP | 02:01+10:01 | 2156 |
| DP | 02:01+11:01 | 3703 |
| DP | 02:01+13:01 | 2046 |
| DP | 02:01+14:01 | 1308 |
| DP | 02:01+17:01 | 4414 |
| DP | 02:02+01:01 | 1664 |
| DP | 02:02+04:01 | 1082 |
| DP | 02:02+05:01 | 1633 |
| DQ | 01:01+02:01 | 7216 |
| DQ | 01:01+02:02 | 1892 |
| DQ | 01:01+03:01 | 6004 |
| DQ | 01:01+03:02 | 3303 |
| DQ | 01:01+05:01 | 45753 |
| DQ | 01:01+05:03 | 6088 |
| DQ | 01:01+06:02 | 7621 |
| DQ | 01:01+06:03 | 5059 |
| DQ | 01:02+02:01 | 8601 |
| DQ | 01:02+02:02 | 2520 |
| DQ | 01:02+03:01 | 5835 |
| DQ | 01:02+03:02 | 2762 |
| DQ | 01:02+03:03 | 2867 |
| DQ | 01:02+04:02 | 1831 |
| DQ | 01:02+05:01 | 8192 |
| DQ | 01:02+05:02 | 6825 |
| DQ | 01:02+06:02 | 37232 |
| DQ | 01:02+06:03 | 6514 |
| DQ | 01:02+06:04 | 10226 |
| DQ | 01:02+06:09 | 2836 |
| DQ | 01:03+02:01 | 5444 |
| DQ | 01:03+03:01 | 3755 |
| DQ | 01:03+03:02 | 1874 |
| DQ | 01:03+05:01 | 5029 |
| DQ | 01:03+06:01 | 3849 |
| DQ | 01:03+06:02 | 7627 |
| DQ | 01:03+06:03 | 18264 |
| DQ | 01:04+05:03 | 3274 |
| DQ | 02:01+02:01 | 13230 |
| DQ | 02:01+02:02 | 11757 |
| DQ | 02:01+03:01 | 9627 |
| DQ | 02:01+03:02 | 2900 |
| DQ | 02:01+03:03 | 10517 |
| DQ | 02:01+05:01 | 6958 |
| DQ | 02:01+06:02 | 9263 |
| DQ | 02:01+06:03 | 4769 |
| DQ | 03:01+02:01 | 4017 |
| DQ | 03:01+03:01 | 7176 |
| DQ | 03:01+03:02 | 20999 |
| DQ | 03:01+03:03 | 1731 |
| DQ | 03:01+05:01 | 5017 |
| DQ | 03:01+06:02 | 3991 |
| DQ | 03:01+06:03 | 2610 |
| DQ | 03:03+02:02 | 868 |
| DQ | 03:03+03:01 | 6771 |
| DQ | 04:01+03:01 | 2001 |
| DQ | 04:01+04:02 | 9455 |
| DQ | 05:01+02:01 | 28919 |
| DQ | 05:01+03:01 | 15260 |
| DQ | 05:01+03:02 | 4454 |
| DQ | 05:01+03:03 | 1827 |
| DQ | 05:01+04:02 | 1582 |
| DQ | 05:01+05:01 | 6692 |
| DQ | 05:01+06:02 | 12109 |
| DQ | 05:01+06:03 | 5679 |
| DQ | 05:01+06:04 | 2696 |
| DQ | 05:05+02:01 | 2202 |
| DQ | 05:05+02:02 | 2975 |
| DQ | 05:05+03:01 | 8176 |
| DQ | 05:05+03:02 | 819 |
| DQ | 05:05+05:01 | 2208 |
| DQ | 05:05+06:02 | 1921 |
| DR | 01:01 | 28642 |
| DR | 01:02 | 8167 |
| DR | 03:01 | 38420 |
| DR | 04:01 | 38453 |
| DR | 04:02 | 3828 |
| DR | 04:03 | 10164 |
| DR | 04:05 | 4203 |
| DR | 07:01 | 51630 |
| DR | 08:01 | 7770 |
| DR | 09:01 | 4672 |
| DR | 10:01 | 4692 |
| DR | 11:01 | 35732 |
| DR | 11:04 | 5429 |
| DR | 12:01 | 5519 |
| DR | 13:01 | 19838 |
| DR | 13:02 | 20913 |
| DR | 13:03 | 9105 |
| DR | 14:01 | 4529 |
| DR | 14:54 | 3106 |
| DR | 15:01 | 51570 |
| DR | 15:02 | 3749 |
| DR | 16:01 | 6856 |
